# Protein-only centromeric chromatin assembly streamlines human artificial chromosome formation

**DOI:** 10.64898/2026.01.28.702397

**Authors:** Gabriel J. Birchak, Praveen Kumar Allu, Prakriti Kashyap, Guhan K. Anbalagan, Shu-Cheng Chuang, Glennis A. Logsdon, Daniel G. Gibson, John I. Glass, Ben E. Black

**Affiliations:** Department of Biochemistry & Biophysics, Perelman School of Medicine, University of Pennsylvania, Philadelphia, PA, 19104 USA; Penn Center for Genome Integrity, Perelman School of Medicine, University of Pennsylvania, Philadelphia, PA, 19104 USA; Epigenetics Institute, Perelman School of Medicine, University of Pennsylvania, Philadelphia, PA, 19104 USA; Graduate Program in Cell & Molecular Biology, Perelman School of Medicine, University of Pennsylvania, Philadelphia, PA, 19104 USA; Department of Genetics, Perelman School of Medicine, University of Pennsylvania, Philadelphia, PA, 19104 USA; J. Craig Venter Institute, La Jolla, CA 92037 USA

**Keywords:** human artificial chromosome, synthetic biology, chromosome biology, centromere, epigenetics, spheroplast fusion

## Abstract

Human artificial chromosomes (HACs) are inherited through cell divisions alongside natural chromosomes, serving as tools for interrogating chromosomal elements and as vectors for large genetic cargoes (*1–5*). Despite recent progress (*6–8*), large (i.e., multiple Mb) HACs have not been reported. Further, HAC formation via epigenetic seeding of centromeric chromatin currently requires prior engineering of recipient human cells (*6–8*), hampering their potential deployment in many useful cell types. Here, we designed, built, and delivered to human cells a 2 Mb HAC construct that is ∼3 times larger than the prior generation. We also report a robust epigenetic centromere seeding approach that initiates immediately upon delivery to the human cell cytoplasm and bypasses genetic engineering of target cells. The HACs are then faithfully inherited in the absence of selection. Thus, formation of functional centromeric chromatin in the same cell cycle of HAC delivery drives high efficiency HAC formation.

**Teaser:** A system for efficient protein-only centromere seeding permits visualizing the first step of HAC formation

## Introduction

Microbial synthetic genomics has succeeded in recoding entire chromosomes and genomes (*9–15*). This, in turn, provides the means to design functionality into biological systems at scales that cannot be approached with even the most expansive gene engineering or gene editing technologies within plausible reach (*16, 17*). The eukaryotic system with the most progress in synthetic genomics is the budding yeast, *Saccharomyces cerevisiae* (*18, 19*). Budding yeast, unlike most eukaryotic species, have small, DNA sequence-defined centromeres (*20*), the loci that drive faithful chromosome inheritance at cell division (*21, 22*). In many eukaryotes, including mammals, functional centromeres are instead defined epigenetically by the presence of a high local concentration of nucleosomes in which histone H3 is replaced with centromere protein A (CENP-A) (*23, 24*). The formation of a functional centromere is essential for generating human artificial chromosomes (HACs). Instead of simply including a DNA sequence-defined centromere element into a HAC construct, HAC formation requires the acquisition of CENP-A-containing chromatin. A general strategy for new centromere formation has been to seed CENP-A nucleosomes at a defined locus (e.g., an array of Lac Operator [LacO] sites) by targeting the CENP-A chaperone, HJURP (e.g., via generating a fusion protein to the Lac Repressor [LacI]) (*25–29*) (fig. S1). Indeed, we have used a pulse of targeted CENP-A nucleosome assembly to form functional centromeres on incoming HAC DNA to efficiently stimulate HAC formation (*6–8*). To date, this and prior HAC formation strategies require selection for antibiotic resistance conferred by the HAC so that by the time HACs are visualized they already contain a functional centromere. Thus, centromere establishment on HACs has not been visualized. After centromere establishment, however, it is clear that no further seeding is necessary, as centromeric chromatin on the HAC is maintained by the same epigenetic process used by natural chromosomes (*30–36*).

The largest HAC construct reported is a 756 kb YAC (*7, 8*). Prior generations of smaller constructs (40-200 kb) undergo uncontrolled and extensive multimerization of the input DNA to generate oligomers typically in the 5-10 Mb range (*1–4, 6*). Uncontrolled multimerization is undesirable because it does not allow one to control the copy number of the genetic cargo delivered to the target cells. The 756 kb YAC is large enough to avoid rampant multimerization (*7, 8*), but it is still not nearly as large (multiple Mb range) as those employed in microbial efforts (*37, 38*). Recent HAC developments have spurred the call to translate microbial synthetic biology triumphs to higher order plant (*39*) and mammalian (*16*) systems. In considering these or any other applications requiring large (>1 Mb) DNA cargo, it remains unclear if any construction or delivery approach for these large cargoes is compatible with HAC technology. Another important current limitation exists in the approach to drive centromere formation via CENP-A nucleosome seeding. This process requires that recipient human cells are pre-engineered to provide a pulse of LacI-HJURP expression (*6–8*) (fig. S1). Avoiding pre-engineering, altogether, would extend HAC formation to “naïve” cells, expediting their utility in diverse cell types with research and clinical significance. In this study, we address both the size limitation and the issues created by the requirement to seed CENP-A nucleosomes. To do this, we built and tested a 2 Mb HAC construct, and we engineered yeast to express and load LacI-HJURP protein onto the HAC construct prior to its delivery. With this approach, we achieve high rates of functional HAC formation without prior genetic manipulation or drug selection of the recipient human cells. We further leverage this efficiency to elucidate early centromere acquisition and nuclear incorporation, shedding light on a process that has evaded scientific inquiry since the first report of epigenetic centromere seeding on HACs.

## Results

### Design and stepwise assembly of a 2 Mb HAC construct

In designing a 2 Mb HAC construct, we considered the “stuffer” DNA provided by prokaryotic genomic sequences. Entire genomes from several prokaryotes have been modified to harbor YAC core elements and then isolated in budding yeast (*40–43*). We chose the ∼1.8 Mb genome from *Haemophilus influenzae* (*Hi*) as a proof of principle for a > 1 Mb genetic cargo for HAC technologies. Our construct design also includes the addition of the functional modules (Fig. 1A) that were in our prior HACs (*7, 8*) that used a synthetic *Mycoplasma mycoides* (*Mm*) minimal genome (JCVI-syn3B (*44*)) as the stuffer DNA. The first step was to add a Cas9 expression plasmid to the yeast strain in order to increase the efficiency of transformation-associated recombination (TAR) cloning, followed by the addition of a mammalian expression cassette encoding tdTomato and NeoR (Fig. 1B, and fig. S2) to screen fusion efficiency and enrich for HAC formation events, respectively. We had planned to use the *TRP1* auxotrophic marker for subsequent TAR cloning but found that strains harboring a YAC with the *Hi* genome surprisingly grew, albeit slowly, on media lacking tryptophan despite the absence of a functional copy of the *S. cerevisiae TRP1* gene (Fig. 1B, and fig. S3). This result is consistent with previous findings that fungal *Trp1* can complement a prokaryotic *trpCF* null mutant (*45*) and that YACs harboring prokaryotic DNA exhibit promiscuous transcription in *S. cerevisiae* (*43*). We hypothesized that ectopic *trpCF* expression was enabling yeast growth in the absence of tryptophan. Consequently, we designed a strategy to remove *trpCF* by replacing it with the *ADE2* marker (fig. S3). Indeed, removal of *trpCF* from the *Hi* YAC resulted in a strain that cannot grow on media lacking tryptophan (Fig. 1B, and fig. S3), permitting selection with *TRP1*. For the final TAR cloning step, we added a ∼10 kb array of LacO and a ∼180 kb region of human chromosome 4 (4q21) that is known to be capable of housing functional centromeric chromatin (*6*) (Fig. 1, B and C).

**Figure 1:**
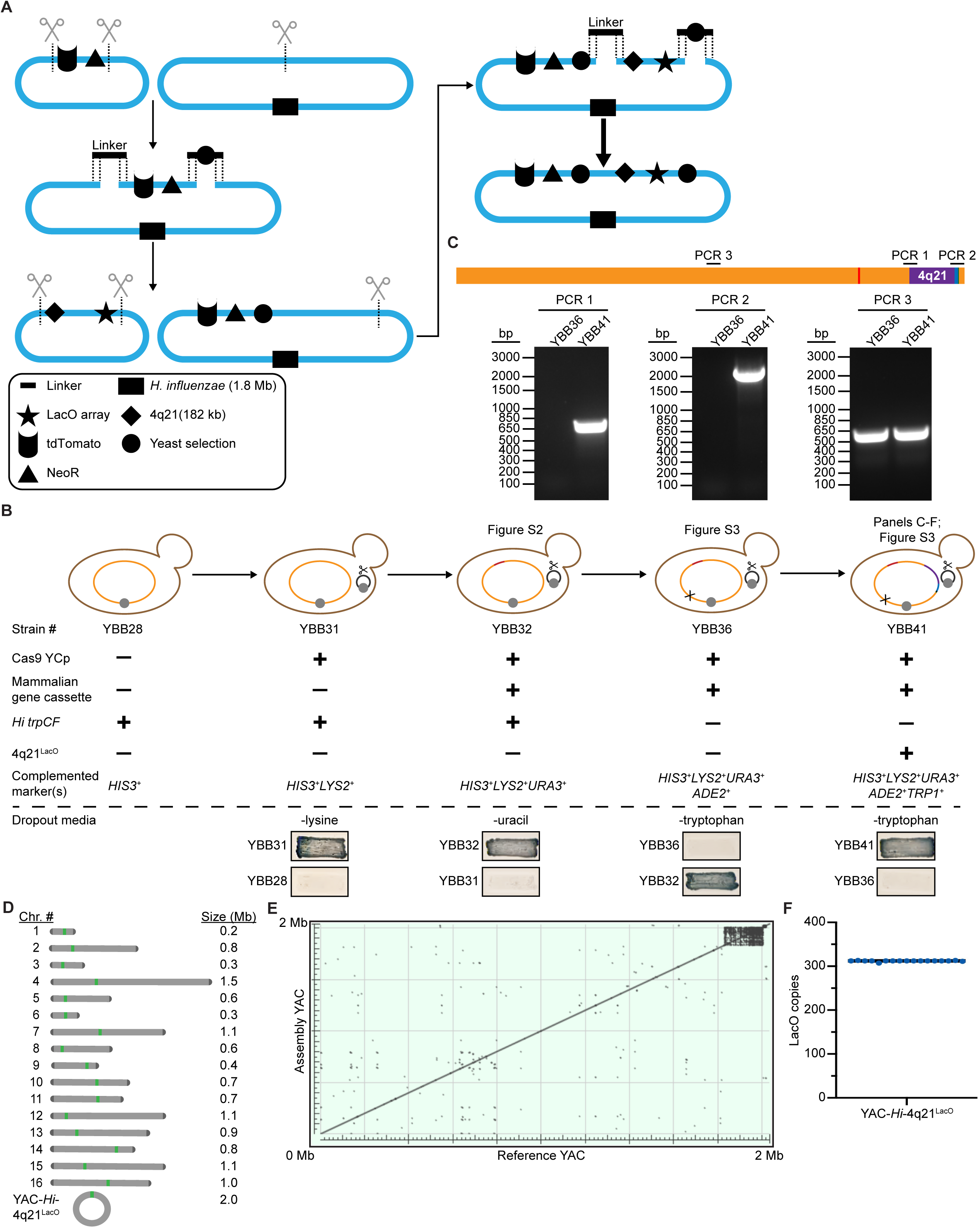
Design, assembly, and long-read ONT sequencing analysis of the yeast strain harboring YAC-*Hi*-4q21^LacO^. (A) Graphical depiction of YAC-*Hi*-4q21^LacO^ engineering approach with all germane genetic elements annotated. (B) Diagrams of serially engineered yeast strains with yeast patches showing auxotrophic status in the indicated dropout conditions. See Table S1 for more details. (C) Colony PCR results from the *H. flu*-4q21 junction (PCR 1), *TRP1*-*H. flu* junction (PCR 2), and stuffer DNA as a control for gDNA quality (PCR 3). (D) Representation of the yeast chromosomes derived from ONT sequencing and an unbiased assembly of pass-filter reads. (E) Dot plot generated by BLAST alignment of the *de novo* assembled YAC-*Hi*-4q21^LacO^, referred to as “Assembly YAC,” against the predicted recombinant YAC, referred to as “Reference YAC.” (F) Number of LacO copies for each read spanning the repetitive array. Bar denotes median LacO copy # = 312 for the 17 reads (individually represented by blue dots) analyzed.

Long-read Oxford Nanopore Technologies (ONT) sequencing and *de novo* assembly of the YBB41 strain harboring YAC-*Hi*-4q21^LacO^ confirmed the overall karyotype (Fig. 1D). Alignment of reads back to our *de novo*-assembled YAC revealed contiguous coverage (fig. S3E) and a dot plot generated via alignment of the *de novo*-assembled YAC against its *in silico* prediction supports a match to the overall design (Fig. 1E). Further, TAR cloning junctions are intact, as revealed by reads spanning them (fig. S3F). The array of LacO repeats is the most challenging region for surviving recombination-based assembly (*46, 47*), as well as the most challenging for sequencing. Using a strategy to identify long reads completely spanning the LacO array, we found the array to be highly homogenous with a median length of 12.43 kb, equivalent to 312 LacO repeats (Fig. 1F). Together, these findings indicate that the derived yeast strain contains all design elements of YAC-*Hi*-4q21^LacO^ and is stably maintained.

### HAC formation with the 2 Mb YAC-*Hi*-4q21^LacO^

To test the ability of YAC-*Hi*-4q21^LacO^ to form functional HACs, we used the same overall strategy we employed previously (*7, 8*) to form HACs with YAC-*Mm*-4q21^LacO^ (Fig. 2A, and fig. S4), inducing LacI-HJURP expression in the recipient cells to promote seeding of CENP-A nucleosomes. We introduced YAC-*Hi*-4q21^LacO^ by spheroplast fusion to human cells expressing dox-inducible mCherry-LacI-HJURP. ∼3 weeks after fusion and antibiotic selection, we screened for HAC formation by immunofluorescence (IF) and fluorescence *in situ* hybridization (FISH) to label CENP-A and the LacO array, respectively. We found that 16.5 +/- 1.2% cells harbor a detectable HAC that is positive for the LacO array and with replicated centromeres, each containing CENP-A (Fig. 2, B and C). We noted, however, that even without induction of LacI-HJURP expression, HACs were detectable at a rate of about half of that observed when LacI-HJURP expression is induced (Fig. 2C, and fig. S4). This finding could be explained by two possibilities: 1) Unlike YAC-*Mm*-4q21^LacO^, YAC-*Hi*-4q21^LacO^ readily forms functional centromeric chromatin independently of epigenetic centromere seeding, or 2) YAC-*Hi*-4q21^LacO^ is very sensitive to low levels of LacI-HJURP, permitting epigenetic centromere seeding via leaky expression of LacI-HJURP in the absence of doxycycline-mediated induction.

**Figure 2:**
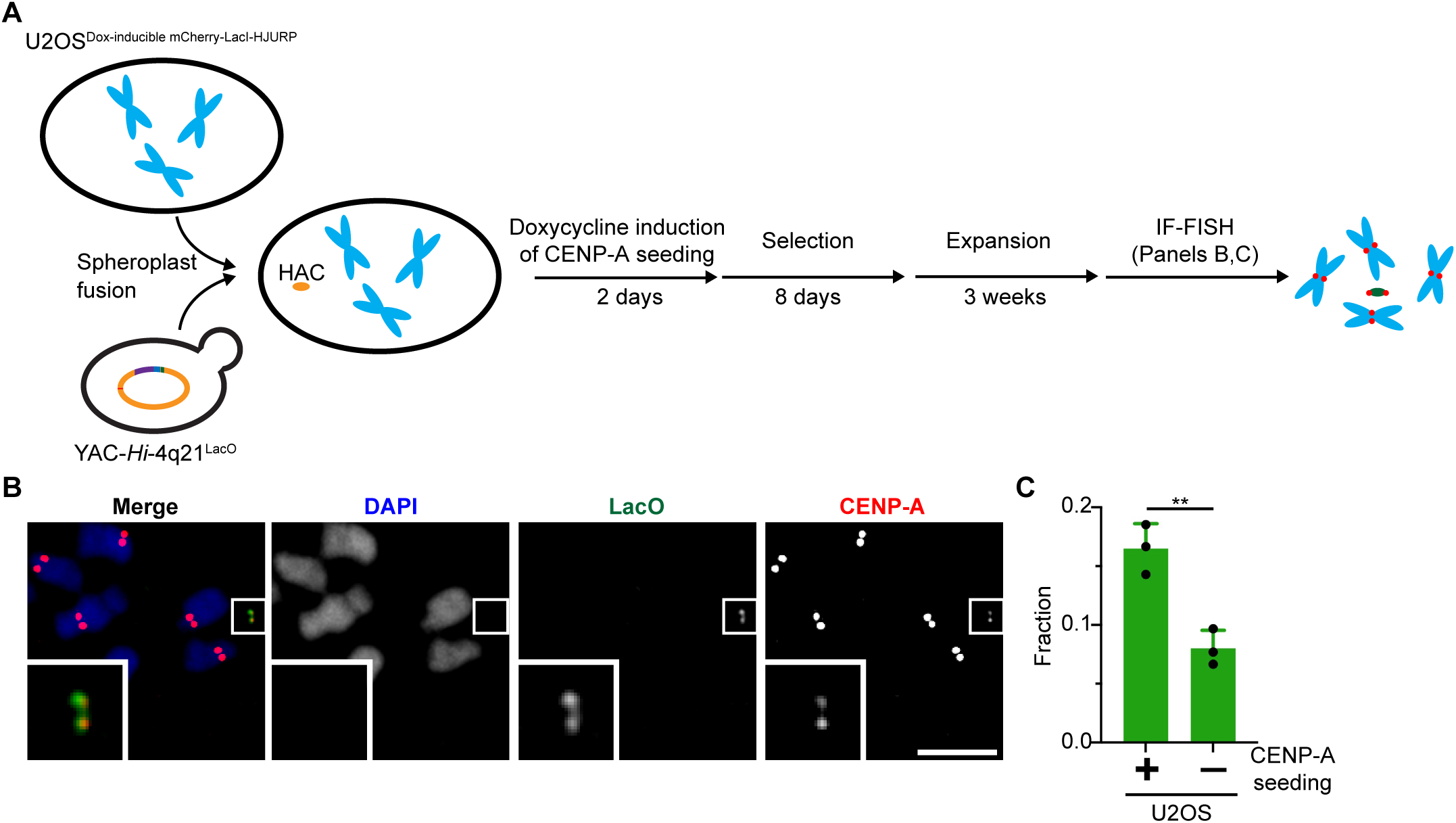
HAC formation with YAC-*Hi*-4q21^LacO^. (A) Graphical depiction of experiment. (B) Representative image of HAC formed in U2OS^Dox-inducible^ ^mCherry-LacI-HJURP^ cells treated with doxycycline following fusion with YAC-*Hi*-4q21^LacO^ spheroplasts. Bar, 5 µm. Inset, 5X magnification. (C) Quantification of HAC formation frequency +/- doxycycline-mediated induction of mCherry-LacI-HJURP expression. The mean (+/- standard error) is shown. The increase in HAC frequency was assessed using an unpaired, one-tailed t test. ** denotes p < 0.01, n=3 (≥ 20 cells/replicate).

### A new one-step approach for forming functional HACs

Given the possibility that even low levels of LacI-HJURP protein may be sufficient to drive efficient formation of centromeric chromatin, we designed a strategy to deliver LacI-HJURP protein along with YAC-*Hi*-4q21^LacO^ by expressing LacI-HJURP in the yeast strain, rather than in the mammalian cell. For this purpose, we integrated a LacI-HJURP expression cassette into a natural yeast chromosome (Fig. 3A). Natural yeast chromosomes lack functional mammalian centromeres, and they are not retained in mammalian cells (*7*). Thus, the LacI-HJURP protein would be present only transiently after spheroplast fusion to seed CENP-A nucleosomes that could establish functional centromeric chromatin that is epigenetically propagated alongside the natural human chromosomes. We installed LacI-HJURP in the yeast strain harboring YAC-*Hi*-4q21^LacO^ at the endogenous *TPI1* locus on yeast chromosome 4, with its expression driven off of the natural *TPI1* gene promoter (Fig. 3A). We isolated a strain with the desired integration and used long-read whole genome ONT sequencing to determine that the desired LacI-HJURP gene addition was achieved and also confirmed that the YAC and other chromosome features, including the LacO array, did not undergo any important changes (Fig. 3, B to E, and fig. S5A).

**Figure 3:**
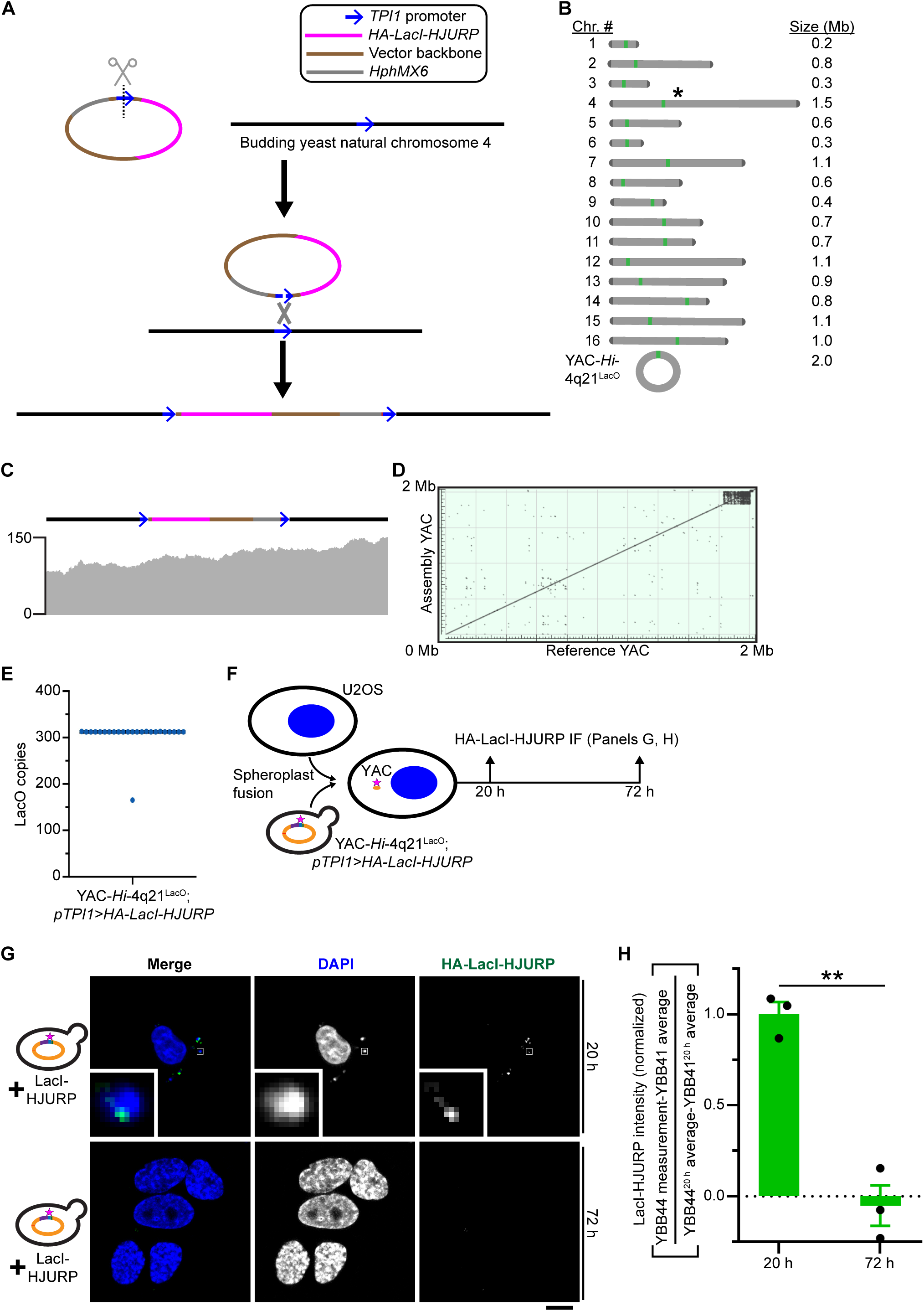
Generating a yeast strain that co-delivers HA-LacI-HJURP protein and YAC-*Hi*-4q21^LacO^. (A) Graphical depiction of yeast integrating plasmid recombining at its target in natural chromosome 4. (B) YAC-*Hi*-4q21LacO; *pTPI1>HA-LacI-HJURP* karyotype verified by ONT sequencing. The asterisk marks the location on natural yeast chromosome 4 of the LacI-HJURP gene expression cassette. (C) The Integrated Genomics Viewer alignment track for the region containing the expected integration along with 5 kb of flanking yeast chromosome 4 DNA revealing contiguous, SNP-free coverage. (D) Dot plot generated by BLAST alignment of the *de novo* assembled YAC-*Hi*-4q21^LacO^, referred to as “Assembly YAC,” against the predicted recombinant YAC, referred to as “Reference YAC.” (E) Number of LacO copies from reads spanning the entire array. Bar denotes median LacO copy # = 312 for the 24 reads (individually represented by blue dots) analyzed. (F) Diagram of approach to measure HA-LacI-HJURP protein delivered from yeast to recipient U2OS cells. (G) Representative images of HA-LacI-HJURP protein detection at the indicated timepoints in U2OS cells fused with YAC-*Hi*-4q21^LacO^; *pTPI1>HA-LacI-HJURP* spheroplasts. Bar, 10 µm. Inset, 10X magnification. (H) Quantification of HA-LacI-HJURP protein per cell. To correct for signal not derived from HA-LacI-HJURP, timepoint-matched averages of U2OS cells fused with YAC-*Hi*-4q21^LacO^ (see fig. S5C) were subtracted from measurements of U2OS cells fused with YAC-*Hi*-4q21^LacO^; *pTPI1>HA-LacI-HJURP* spheroplasts. All data are normalized to the difference in the average signal value per cell of U2OS cells fused with YAC-*Hi*-4q21^LacO^; *pTPI1>HA-LacI-HJURP* spheroplasts and U2OS cells fused with YAC-*Hi*-4q21^LacO^ at 20 h (see **Materials and Methods** for further details). The difference in LacI-HJURP was analyzed using an unpaired, two-tailed Welch’s t test. ** denotes p < 0.01, n=3 (≥ 22 cells/replicate).

We first monitored the presence and localization of the HA-tagged LacI-HJURP fusion protein upon delivery to “naïve” U2OS cells (Fig. 3F). At an early timepoint (20 h), HA-LacI-HJURP that had been expressed in the yeast cells prior to spheroplast fusion is detectable as foci in the cytoplasm (Fig. 3G). We note that the cytoplasmic HA-LacI-HJURP foci typically overlap with a portion of larger DAPI-positive foci that likely represent budding yeast nuclei. This localization pattern is consistent with HA-LacI-HJURP being present on the HAC construct at an early timepoint. At a later timepoint (72 h), the HA-LacI-HJURP is not detectable at any location in the cells (Fig. 3, G and H). Note, that nearly all U2OS cells were initially fused with the yeast strain containing HA-LacI-HJURP protein and YAC-*Hi*-4q21^LacO^ (fig. S5). We conclude that HA-LacI-HJURP is in position to assemble CENP-A-containing nucleosomes, and that, as we anticipated, it is degraded and diluted as cells grow and divide.

Since HA-LacI-HJURP and HAC DNA are present together in the cytoplasm where nascent CENP-A (and its partner core histones) are expressed, we reasoned that the formation of functional centromeric nucleosomes might initiate in the cytoplasm. We focused on endogenous CENP-C (Fig. 4A), an essential centromere component (*48*), since it recognizes CENP-A only in the context of a fully assembled nucleosome, with direct recognition of H2A/H2B heterodimers and the (CENP-A/H4)_2_ heterotetramer on the surface of the nucleosome (*49, 50*). At an early timepoint, both strains with and without LacI-HJURP deliver YAC-*Hi*-4q21^LacO^ (detected via FISH against LacO) to the U2OS cell cytoplasm (Fig. 4B). Colocalization of YAC-*Hi*-4q21^LacO^ and CENP-C was detectable only when using the strain expressing LacI-HJURP (Fig. 4, B and C). After cells grow and divide to a later (72 h) timepoint, essentially no cells harboring YAC-*Hi*-4q21^LacO^ are detected for fusions with the strain lacking LacI-HJURP (Fig. 4, B and C), consistent with their degradation if they do not rapidly form functional centromeres. By contrast, for the strain expressing LacI-HJURP, the localization of YAC-*Hi*-4q21^LacO^ moves from the cytoplasm to the nucleus where they remain colocalized with CENP-C (Fig. 4, B and C). We conclude that functional centromeric chromatin formation initiates in the cytoplasm, and that this endows the HACs to join the natural human chromosomes at the time of cell division.

**Figure 4:**
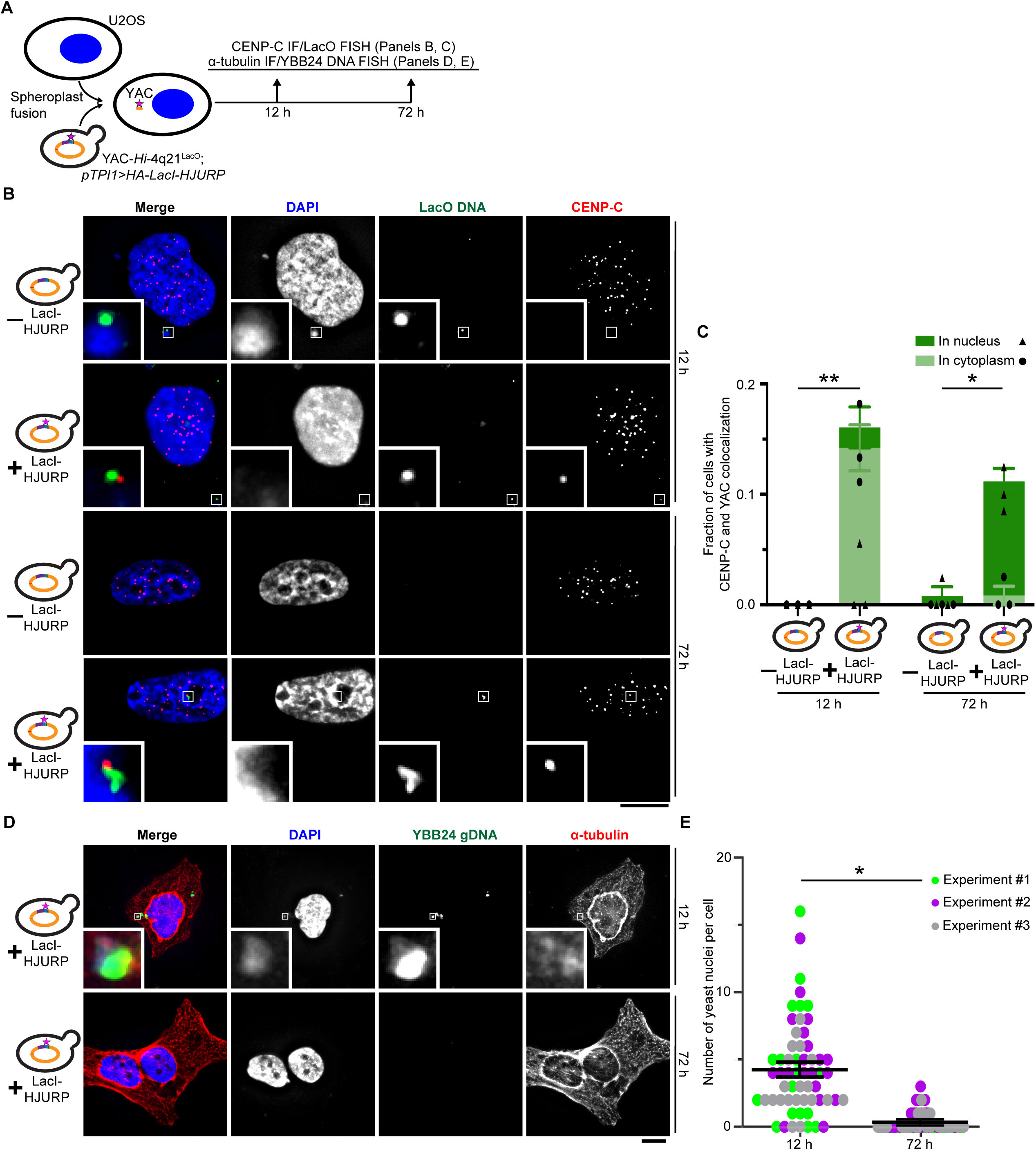
Protein-only centromere seeding is initiated in the cytoplasm, conferring efficient nuclear incorporation of YAC-*Hi*-4q21^LacO^. (A) Diagram of approach to assess acquisition of CENP-C on YAC-*Hi*-4q21^LacO^ or persistence of yeast genomic material at the indicated timepoints. (B) Representative images of U2OS cells from fusions with either the YAC-*Hi*-4q21^LacO^ or YAC-*Hi*-4q21^LacO^; *pTPI1>HA-LacI-HJURP* yeast strains and processed for CENP-C IF, LacO FISH 12 h or 72 h post fusion. Bar, 10 µm. Inset, 6X magnification. (C) Quantification of coincident LacO FISH and endogenous CENP-C in either the nuclear or cytoplasmic compartments of the cell at the indicated timepoints after spheroplast fusion. The increase in the fraction of cells carrying a CENP-C+ YAC from fusions with the YAC-*Hi*-4q21^LacO^; *pTPI1>HA-LacI-HJURP* strain relative to the YAC-*Hi*-4q21^LacO^ strain, regardless of compartment location, was analyzed using one-tailed, unpaired Welch’s t tests. * denotes p < 0.05, ** denotes p < 0.01, n=3 (≥ 10 cells/replicate). (D) Representative images of U2OS cells fused to the YAC-*Hi*-4q21^LacO^; *pTPI1>HA-LacI-HJURP* yeast strain and cultured for the indicated timepoints before conducting ⍺-tubulin IF and YBB24 gDNA FISH. Bar, 10 µm. Inset, 10X magnification. (E) Quantification of yeast nuclei in U2OS cells. The difference in yeast nuclei between timepoints was analyzed using a two-tailed, unpaired Welch’s T test. * denotes p < 0.05, n=3 (≥19 cells/replicate). Each cell constituting the measurements is plotted and colored according to the independent experiment performed. The average of experiment means +/- standard error of experiment means is overlaid. Note that 10.4 +/- 0.01% of U2OS cells did not have any yeast nuclei at 12 h; whereas, 73.2 +/- 15.4% of U2OS cells did not have any yeast nuclei at 72 h.

Natural yeast chromosomes, different from the HAC construct, lack functional human centromeres or LacO sites, thus we anticipated that, like YAC-*Hi*-4q21^LacO^ coming from strains lacking LacI-HJURP protein (Fig. 4, B and C), natural yeast chromosomes would be lost through cell divisions following their co-delivery via our spheroplast fusion approach. To test this, we detected yeast chromosomes via FISH using genomic DNA from the overall parental yeast strain (Fig. 4, D and E; YBB24; i.e. a strain without any part of the HAC construct present). At an early timepoint after spheroplast fusion (12 h), we found 4.8 +/- 0.6 yeast nuclei within fused U2OS cells (Fig. 4, D and E). At a later timepoint (72 h), we observed a nearly complete loss of yeast nuclei (0.4 +/- 0.2; Fig. 4, D and E). We conclude that yeast chromosomes lacking a functional human centromere remain in the human cell cytoplasm and are degraded within a few cell cycles after spheroplast fusion.

Given our success with U2OS cells in achieving highly efficient cell fusions and the ability to observe early centromere formation events in cells without selection, we designed a modified scheme to detect functional HACs in the absence of any selection whatsoever (Fig. 5A). We also omitted mitotic arrest, which had been previously used based on earlier experiments to increase random YAC inclusion in nuclei at mitotic exit (*51*), since early cytoplasmic centromere formation steps appear to lead to efficient nuclear inclusion of HACs (Fig. 4, B and C). With this modified approach, we found HACs in 14.5 +/-2.7% cells; whereas almost no HACs (1.2 +/- 1.2% cells) were detected with the precursor yeast strain that harbors YAC-*Hi*-4q21^LacO^ but lacks LacI-HJURP expression (Fig. 5, B and C). We conclude that HACs formed in the absence of induced gene expression of LacI-HJURP in the pre-engineered recipient cells (Fig. 2, and fig. S4) were a result of sensitivity to low levels of LacI-HJURP. As with the prior strategy (*7, 8*) (Fig. 2), the HACs that formed were uniformly small (Fig. 5B) in appearance with no sign of the rampant multimerization exhibited by smaller BAC-derived HACs (*3–6*). Our experiments, in contrast to all prior HAC experiments of which we are aware, demonstrate that HACs can be readily identified in cell populations without selection or other means of enrichment, streamlining their use.

**Figure 5:**
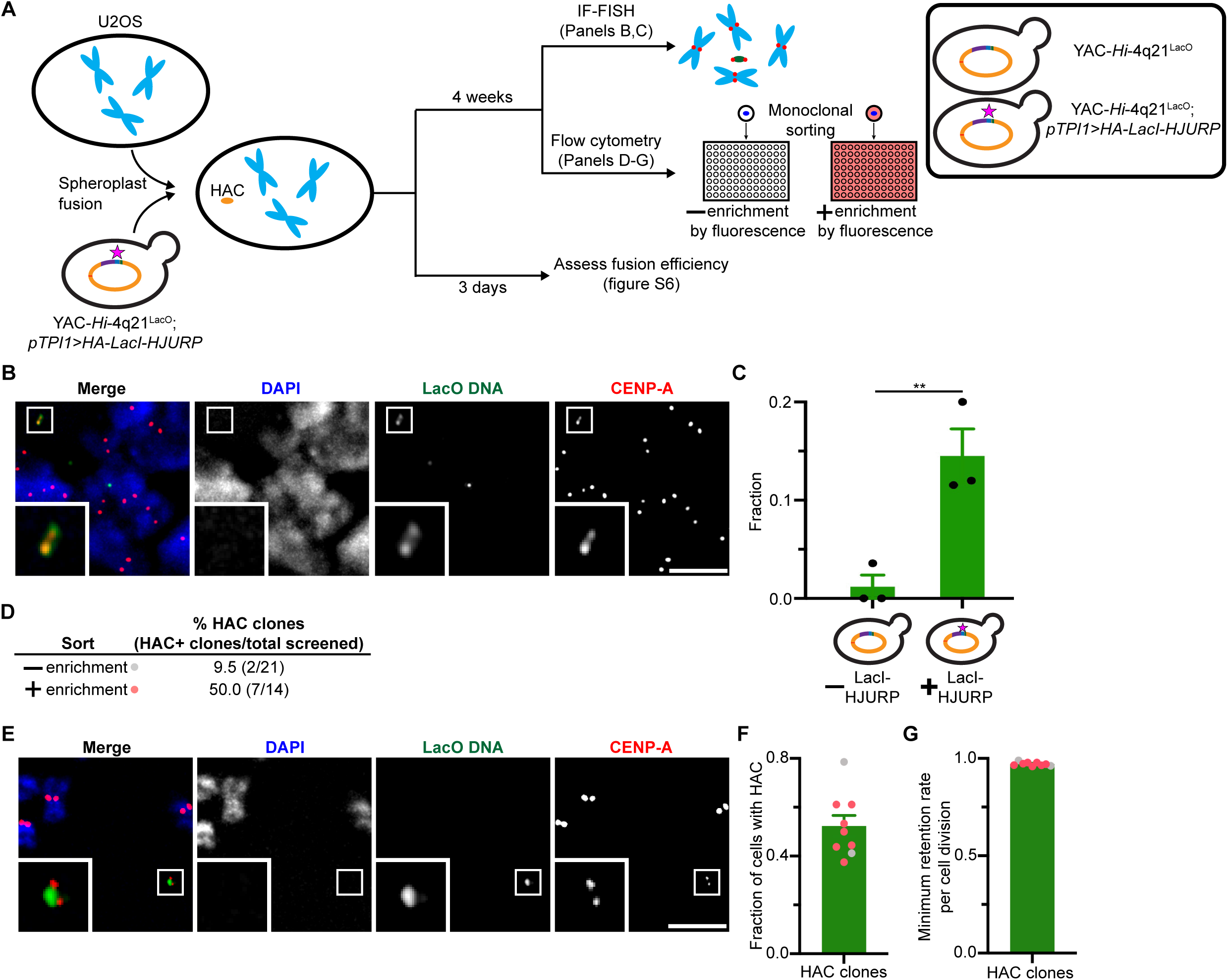
Protein-only centromere seeding efficiently forms HACs and bypasses selection. (A) Outline of the modified HAC formation assay. 3 x 10^5^ U2OS cells were fused with 9 x 10^7^ spheroplasts before splitting two ways to assess fusion efficiency or to subculture and expand for either CENP-A IF-LacO FISH to quantify HAC formation frequency or flow cytometry to derive monoclonal HAC lines (see fig. S7 for gating and controls). (B) Representative image of a HAC generated with the protein-only centromere seeding HAC formation approach using YAC-*Hi*-4q21^LacO^ derived from the YAC-*Hi*-4q21^LacO^; *pTPI1>HA-LacI-HJURP* yeast strain. Bar: 5 µm. Inset, 5X magnification. (C) Frequency of HAC formation. The increase in HAC frequency was assessed using an unpaired, one-tailed t test. ** denotes p < 0.01, n=3 (≥ 25 cells/replicate). (D) Clonal cell lines were grown up to a 6-cm plate before performing IF-FISH to screen for HACs. The fraction of screened monoclonal cell lines harboring HACs is shown. The difference in HAC clones as a fraction of total screened clones was analyzed using a two-sided Fisher’s exact test, p = 0.02. (E) Representative image of a HAC observed in a monoclonal HAC line derived from the sort to enrich for HACs by fluorescence. Bar, 5 µm. Inset, 5X magnification. (F) Graph depicting the fraction of cells in a HAC clone carrying a HAC with individual clones colored according to sorting strategy. (G) Graph depicting the minimum HAC retention rate per cell division, assuming ∼23 cell doublings to reach IF-FISH (see **Materials and Methods**), with individual clones colored according to sorting strategy.

To address stability of HACs in cell culture, we sorted single cells from the non-selection HAC formation assay into individual wells of a 96-well plate, either with or without sorting for red fluorescence generated by the tdTomato cassette present on the HAC (Fig. 5A, and fig. S7). Cell populations arising from single cells were generated and then screened for the presence of HACs. Whereas 9.5% of screened clones derived from the fluorescence-agnostic sort proved to be HAC clones, 50% of screened clones derived from the sort on red fluorescence were HAC positive (Fig. 5, D and E). Populations of cells (after 23+ cell generations; see **Materials and Methods**) arising from a single cell harboring a HAC were found to contain HACs in 35-80% of cells, depending on clonal isolates (Fig 5F). This translates to a minimum retention rate per cell division of 96-99% (Fig. 5G). Therefore, in the absence of selection, the observed HACs present evidence of stability through mitotic cell divisions as a result of centromeres formed via “protein-only” CENP-A nucleosome seeding. Altogether, our experiments indicate that efficient HAC formation is almost entirely dependent upon initial CENP-A nucleosome seeding by the LacI-HJURP fusion protein, resulting in a new centromere on the HAC construct that is subsequently epigenetically propagated as HAC-containing cells grow and divide.

## Discussion

In this study, we have advanced HAC technology in two main areas. **First**, we developed a 2 Mb-sized HAC construct and demonstrated that the spheroplast fusion delivery strategy successfully transmits them to recipient human cells. One could envision translating our HAC construct assembly method to add useful genetic cargo for essentially any rational purpose (Fig. 6A). We speculate that the upper limit of HAC construct design in our system would extend beyond the largest YAC (2.3 Mb) of which we are aware (*52*), perhaps beyond a large engineered yeast chromosome (e.g., 11.8 Mb) (*53*). Megabase-scale HAC design and assembly should now permit the large and complex genetic engineering that has to date been mostly limited to microbial systems. **Second**, “protein-only” epigenetic centromere seeding streamlines the initial approach, allowing HACs to be formed in naïve cells without otherwise manipulating the genome of those cells (Fig. 6B). Since the protein (LacI-HJURP) encoded in yeast is expressed prior to spheroplast fusion, it is subjected to a limited lifetime as a result of degradation and dilution through cell division (Fig. 3, G and H).

**Figure 6:**
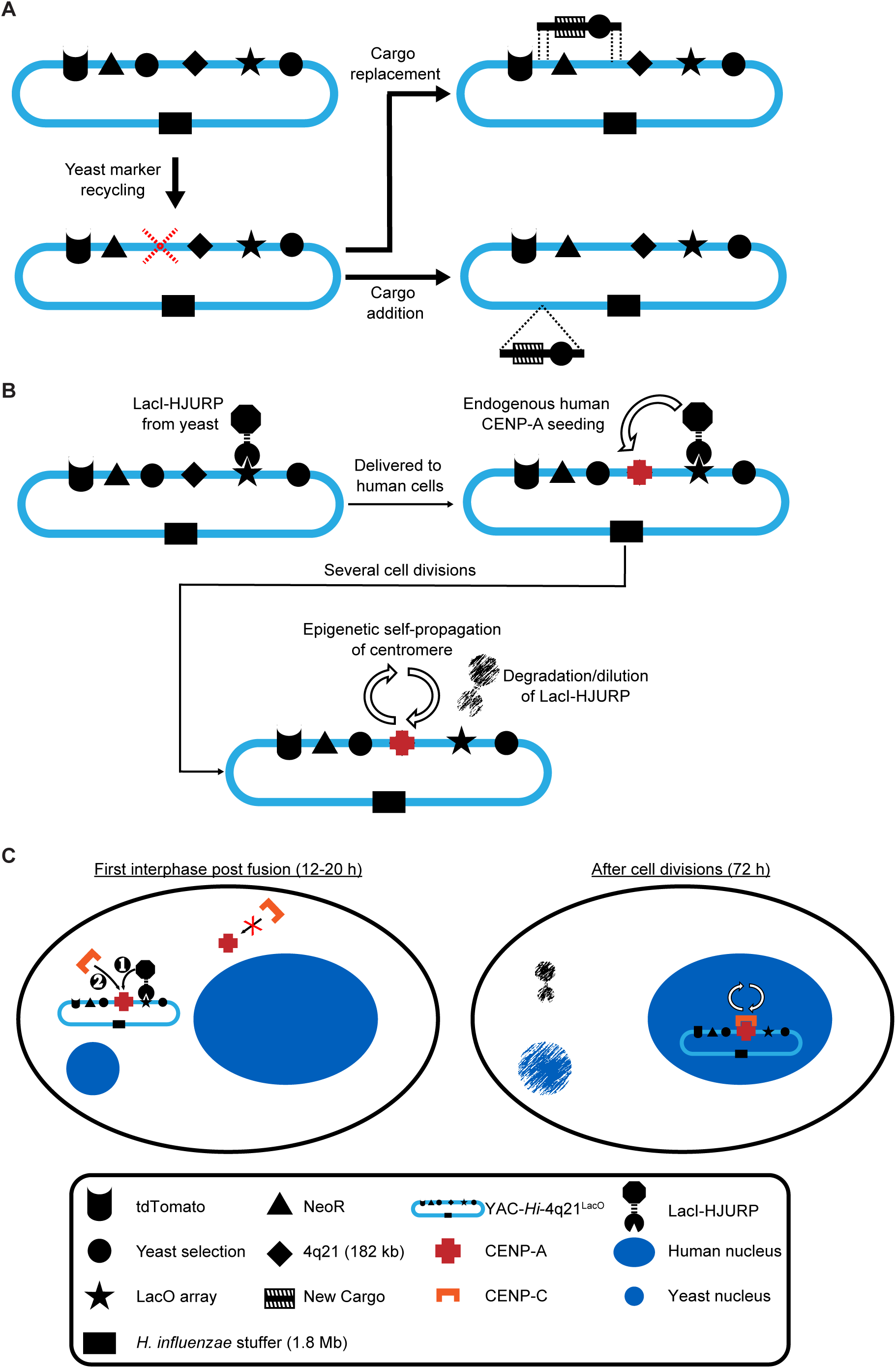
A system for streamlined future HAC development using protein-only centromere seeding. (A) Diagram of how other relevant cargoes could be added to *Hi* YAC using TAR. (B) Diagram of our protein-only seeding strategy and how epigenetic centromere propagation persists even as LacI-HJURP protein is degraded and/or diluted through cell divisions. (C) Diagram of early HAC formation events. Once the YAC and LacI-HJURP protein are delivered to the cytoplasm of the recipient cell, HJURP deposits CENP-A (and H4) and nucleosomes (consisting of two copies each of H2A, H2B, CENP-A, and H4) are established (step 1). This then allows CENP-C, which directly binds nucleosomal CENP-A, to be recruited to the nascent centromere on the HAC (step 2). Following mitosis, the HAC is incorporated in the nucleus where its centromere now persists by the natural epigenetic centromere propagation loop, allowing the molecule to stably persist over many cell divisions while LacI-HJURP protein and the accompanying natural yeast genome are degraded and/or diluted.

Using the streamlined delivery and seeding approach, we revealed that centromere formation on the HAC construct is initiated in the cytoplasm. The source of endogenous human centromeric chromatin components are likely to be nascent polypeptides prior to when they would normally undergo cell cycle-coupled assembly on natural centromeres in the nucleus (*21, 22*). A substantial fraction of the nascent HACs are then incorporated into the nucleus (Fig. 4, B and C). We speculate that this occurs at the exit of mitosis via centromere connections of the HAC to the mitotic spindle (Fig. 6C). To our knowledge, our experiments (Figs. 3, F to H, and 4, A to C) are the first to visualize molecular events during the genesis of a centromere. The original generation of HACs formed functional centromeres in as few as 9 per 2 x 10^6^ cells (*1*), necessitating drug selection to enrich for clonal populations harboring HACs before centromere function could be assessed (*1–3, 5, 6*). With the protein-only epigenetic centromere seeding approach reported here, HACs form in ∼1 out of 7 cells, permitting observation of centromere initiation which occurs before drug selection would ever exhibit an effect.

We also highlight the ability of our streamlined HAC formation approach (Fig. 5A) to bypass mitotic arrest prior to fusion, which had previously been shown to increase the nuclear delivery frequency of YACs without functional human centromeres (*51*). Indeed, in the absence of mitotic arrest, functional HACs formed and were detected without imposing any selective pressure (Fig. 5, A to C). To our knowledge, this is the first report of HACs cytologically detected in the absence of drug selection. If the incoming HAC construct lacks a functional centromere when the human cell enters mitosis, we envision that it would only have a small and random chance of being incorporated into the cell nucleus at mitotic exit since the nucleus is only ∼10% of the cell volume. The presence of a functional centromere, on the other hand, biases inheritance via spindle attachment, and we conclude that this feature is what drives the high efficiency of HAC formation that we observe (Fig. 5, B and C).

For future HAC construct design, the choice of DNA sequence for HAC stuffer DNA does not appear to have a strong requirement for very AT-rich sequences. In our previous report (*7*) describing YAC-*Mm*-4q21^LacO^ where the stuffer DNA has an AT content of 77%, we speculated that the extreme AT-richness may positively contribute to centromere function. YAC-*Hi*-4q21^LacO^ contains a stuffer DNA that is considerably less AT-rich at 62%. This is quite similar to the 59% AT content of the human genome and to human centromeric DNA, which is ∼63-67% (*54*). Thus, at least in the context of a 2 Mb construct, the extreme AT-richness of our previous HAC is unnecessary for centromere formation. We note that instead of multimerized BAC-based HACs with many LacO array foci, both YAC-*Mm*-4q21^LacO^ and YAC-*Hi*-4q21^LacO^ have a “double-dot” or similar appearance on mitotic chromosomes (*7*) (Fig. 2B, 5, B and E, and fig. S4B). This is consistent with YAC-*Mm*-4q21^LacO^ and YAC-*Hi*-4q21^LacO^ each forming functional centromeres without multimerizing.

Looking towards the future, our HAC strategy could be developed for *ex vivo* therapies (e.g., chimeric antigen receptor (CAR) therapies (*55, 56*), hematopoietic stem cell (HSC) correction (*57*), and cell transplantation for degenerative disease treatment (*58, 59*)) that presently employ integrating viruses and endonucleases that impact natural human chromosomes. Bypassing cell line engineering and drug selection to form HACs in a large fraction of cells, should, in principle, limit therapeutic manufacturing time and subculture for *ex vivo* applications. Further, bypassing mitotic arrest should render formation in these non-tumorigenic cell types, which would be susceptible to p53-mediated cell cycle arrest and/or death (*60, 61*), feasible. Finally, expeditious enrichment of HAC populations by flow cytometry will augment efficient HAC formation to bestow polyclonal cell populations useful for therapeutics or biologics production. In the process of developing HACs for these or other specific applications, our HAC technology will be of outstanding research value to recapitulate and interrogate fundamental chromosomal processes.

## Acknowledgements

We thank our UPenn colleagues K. Murakami for discussions and M. Bartolomei, A. Dananberg, and M. Lampson for comments on the manuscript. We also thank E. Makeyev (King’s College) for reagents. The content is solely the responsibility of the authors and does not necessarily represent the official views of the National Institutes of Health. This manuscript is the result of funding in whole by the National Institutes of Health (NIH). It is subject to the NIH Public Access Policy. Through acceptance of this federal funding, NIH has been given a right to make this manuscript publicly available in PubMed Central upon the Official Date of Publication, as defined by NIH.

## Author contributions

G.J.B., J.I.G., and B.E.B. conceived the project. G.J.B., P.K.A., G.A.L., D.G.G., and B.E.B. designed experiments. G.J.B., P.K.A., G.K.A., and P.K. performed experiments. G.J.B., P.K.A., P.K., G.K.A., S.-C.C., G.A.L., and B.E.B. analyzed data. G.J.B. and P.K.A. generated reagents. G.J.B. and B.E.B. wrote the paper. All authors edited the paper.

## Data and materials availability

Sequencing data will be released on Zenodo at the time of publication. All other data required to evaluate the conclusions are present in the paper or supplementary material. No new code was generated for analysis of the data reported in this publication. Reagents used are available from commercial sources or from the corresponding author upon reasonable request.

## Materials and Methods

### gRNA synthesis

Lyophilized sense and antisense IDT DNA Ultramers (Table S2) were resuspended in annealing buffer (10 mM Tris-Cl, 0.1 mM EDTA, 50 mM NaCl, pH 8.0) to 50 µM. Equal volumes of the Ultramers were mixed in a 1.5 ml Eppendorf tube and incubated for 5 min in a heat block set to 95 ℃. The heat block was shut off, and the tubes were left to anneal for 16 h.

sgRNA was then generated by *in vitro* transcription using the MEGAshortscript^TM^ T7 Transcription Kit (Invitrogen catalog #: AM1354) according to manufacturer instructions. In brief, 25 pmol annealed Ultramer was used as template for transcription for 4 h at 37 ℃. 1 µl Turbo DNase was then added and the reaction incubated at 37 ℃ for an additional 15 min to terminate the reaction. The MEGAclear^TM^ Transcription Clean-Up Kit (Invitrogen catalog #: AM1908) was used to purify the transcribed sgRNA, involving two 50 µl elutions with 70 ℃ 10 mM Tris-Cl (pH 7.5).

### TAR cloning

TAR cloning to generate yeast strains YBB32 and YBB41 deployed the spheroplast transformation largely as reported (*62*). In brief, 50 ml of synthetic minimal media with the appropriate dropout was inoculated with a colony and grown overnight to an OD_600_ of 1.0. The culture was pelleted at 1600 x g for 3 min, washed with 50 ml sterile water, pelleted again, and resuspended in 20 ml 1 M sorbitol. The yeast were incubated on ice at 4 °C overnight. The next day, cells were harvested at 1741 x g for 3 min and resuspended in 20 ml SPE (1 M sorbitol, 7.89 mM Na_2_HPO_4_, 2.33 mM NaH_2_PO_4_, 10 mM EDTA pH 8.0). 40 µl of β-mercaptoethanol and 40 µl 10 mg/ml zymolyase-20T (MP Biomedicals catalog #: 320921) were added to the suspension. The tube was inverted several times and then placed in a shaker to incubate at 30 ℃, 50 rpm. After 20 min, the tube was inverted several times and again returned to the shaker to incubate for 20 min. 30 ml of 1 M sorbitol was added to the suspension before harvesting spheroplasts at 1600 x g for 5 min. Spheroplasts were washed with 50 ml 1 M sorbitol before harvesting at 1600 x g for 5 min. Spheroplasts were resuspended in 2.5 ml STC (1 M sorbitol, 10 mM Tris-Cl pH 7.5, 10 mM CaCl_2_) and incubated at room temperature for 10 min. 200 µl of the suspension was added to an Eppendorf tube containing the DNA (see Table S1) and sgRNA along with water to a total volume of 40 µl, and the suspension was incubated for 10 min at room temperature. 1 ml of PEG/CaCl_2_ (20% w/v PEG8000 (Sigma catalog #: 89510), 10 mM CaCl_2_, 10 mM Tris-Cl pH 7.5) was added and the solution mixed by inverting and flicking the Eppendorf tube. The solution was incubated at room temperature for 20 min. Transformed spheroplasts were harvested at 1600 x g for 8 min. The spheroplasts were suspended in 800 µl SOS medium (1 M sorbitol, 6.5 mM CaCl_2_, 0.25% w/v bacto yeast extract, 0.5% w/v bacto peptone) and incubated at 30 ℃ for 1 h. The 800 µl solution was then pipetted onto 8 ml of 55 ℃ synthetic minimal top agar devoid of the appropriate nutrients (YBB32: - histidine-uracil-lysine, YBB41: -histidine-uracil-lysine-tryptophan), inverted 3 times, and poured onto synthetic minimal sorbitol plates for recovery of transformants. Prototrophs were patched onto synthetic minimal agar plates with the matching dropout before genotyping as described in “Genotyping yeast strains” (see Table S2 for primers).

### *trpCF* allele swap

The allele swap was performed using the Markerless Yeast Localization and Overexpression approach, largely as previously reported (*63*). To generate the gRNA, 70-nucleotide primers (Table S2) were used in a PCR reaction, yielding a double-stranded gRNA spacer sequence flanked by 50-nucleotide homology arms. The pCAS vector carrying the *Cas9* expression cassette, *HphMX6* selection gene, and gRNA stuffer was prepared by digesting the vector with BsaI and NotI and purifying with the QIAquick PCR Purification kit (Qiagen catalog #: 28104). *pURA3>ADE2* flanked by 300-bp arms homologous to regions flanking *trpCF* was ordered as a gene block from Twist Biosciences. To avoid unwanted recombination events at the natural *S. cerevisiae ADE2* locus, *ADE2* was codon optimized to eliminate identical stretches ≥ 20 bp.

Transformation was carried out as previously described (*64*) with modifications. YBB32 yeast were grown in 50 ml synthetic minimal (-histidine -uracil -lysine) media to an OD_600_ = 1.0. 2 ml of culture/transformation was pelleted at 4,500 x g, 5 min, room temperature. Cells were resuspended in 1 ml H_2_O and transferred to a microcentrifuge tube before pelleting at 2,400 x g for 1 min. Cells were washed one time in 1 ml 100 mM LiOAc and pelleted. Cells were resuspended in a transformation mix containing 100 µl 50% PEG3350 (Sigma catalog #: 88276), 5 µl 3 M LiOAc, and 10 µl 10 mg/ml boiled salmon sperm DNA per transformation. ∼120 µl of cells and transformation mix were then aliquoted into an Eppendorf tube containing 5 µl of the crude gRNA PCR reaction; 500 ng of the PCR-purified, digested pCAS vector; 1 µg of the *pURA3>ADE2* donor template; and H_2_O (added such that volume of DNA mix = 40 µl). The cells were incubated at 30 ℃ for 30 min, then transferred to 42 ℃ for 30 min, before pelleting again at 2,400 x g, 1 min. Cells were resuspended in 1 ml YPD and recovered for 16 h at 30 ℃. Cells were then plated on synthetic minimal (-histidine-uracil-lysine+ 5 mg/ml adenine plates) containing 1 mg/ml hygromycin B (Invitrogen catalog #: 10687010) and allowed to form colonies. White, drug-resistant colonies were patched on equivalent plates and genotyped as described in “Genotyping yeast strains”, below (see Table S2 for primers).

Recombinant yeast strains were struck out on synthetic minimal (-histidine -uracil -lysine) plates and colonies were replica plated on synthetic minimal (-histidine -uracil -lysine + 1 mg/ml hygromycin B) to identify colonies that had mis-segregated the pCAS vector. YBB36 was then characterized as described in “Yeast strain sequencing, genome assembly, and analysis”, below.

### Introducing the LacI-HJURP expression cassette

A plasmid carrying the *pTPI1>HA-LacI-HJURP* and *HphMX6* expression cassettes was digested with PacI for 3 h. The digestion was run on a 0.5% agarose gel and the linearized product extracted using the Qiaquick Gel Extraction Kit (Qiagen catalog #: 28704). 100 ng of linearized product was electroporated into yeast as reported (*65*). YBB41 transformants were allowed to recover for 16 h and were plated on synthetic minimal (-histidine -uracil -tryptophan -lysine + 1 mg/ml hygromycin B) before patching and genotyping as described in “Genotyping yeast strains” (see Table S2 for primers). Junction-positive candidates were then characterized as described in “Yeast strain sequencing, genome assembly, and analysis,” yielding strain YBB44.

### Genotyping yeast strains

gDNA was extracted from patches of candidate yeast strains as described (*9*) using the QIAprep Spin Miniprep Kit (Qiagen catalog #: 27104). 1 µl of eluted gDNA was used as template for PCRs.

### Yeast strain sequencing, genome assembly, and analysis

YBB32 and YBB36 yeast strains were grown in 150 ml appropriate synthetic minimal media (-histidine-uracil-lysine) before harvesting. Specifically, a 5 ml yeast starter was used to inoculate 145 ml media, which was then grown overnight in a shaking incubator set to 30 ℃, 183 rpm. When OD_600_ = 0.5, the yeast were distributed to three 50 ml conical tubes before spinning down at 700 x g, 16 ℃, 15 min. Each pellet was resuspended in 10 ml PBS (4 ℃) before pooling into a single tube. The yeast were again harvested at 700 x g, 16 ℃, 15 min. Cells were washed in 30 ml PBS (4 ℃) and again harvested at 700 x g, 16 ℃, 15 min. The pellet was resuspended in 20 ml cold SPE media. 40 µl of β-mercaptoethanol and 200 µl 10 mg/ml zymolyase-20T were added, the tube was gently inverted to mix, and then placed in a shaking incubator set to 30 ℃, 25 rpm for 1 h. After 1 h, 100 µl of yeast solution was added to either 1 M sorbitol or 2% SDS and the OD_600_ values measured. If the ratio of the measurement in sorbitol versus that in SDS was greater than 5:1, the spheroplasts were harvested at 700 x g, 16 ℃, 15 min. The pellet was then resuspended in 150 µl cold 500 mM NaCl and transferred to a 2 ml Eppendorf tube. From here, high molecular weight gDNA was prepared using the Monarch HMW DNA Extraction Kit for Tissue (NEB catalog #: T3060) according to the manufacturer’s instructions; albeit, isopropanol precipitation and centrifugation were used to purify and clean the gDNA rather than utilizing the included beads.

Short fragments were removed from the gDNA using the SRE XS kit (PacBio catalog #: 102-208-200). The pellet of gDNA was resuspended in 100 µl of included LTE buffer. Depletion of short fragments was validated with the Agilent TapeStation 4200.

Sequencing libraries were prepared using 1 µg of input gDNA and the Native Barcoding Kit 24 V14 (Oxford Nanopore Technologies catalog #: SQK-NBD114.24), and washes were carried out with short fragment buffer. The entirety of the sample library was mixed with library beads and loaded onto an R10.4.1 flow cell and sequenced on a PromethION device.

Plasmidsaurus was commissioned to sequence YBB41 and YBB44. 50 ml of synthetic minimal media (-histidine-uracil-lysine-tryptophan) was inoculated with the appropriate strain and grown to OD_600_ = 1.0. Yeast cells were pelleted at 500 x g for 2 min. The pellet was resuspended in 0.5 ml of DNA/RNA shield (Zymo Research catalog #: R1100) before sending to Plasmidsaurus for ONT sequencing. In the case of YBB44, sufficient LacO coverage was obtained in a single sequencing attempt. In the case of YBB41, sequencing was carried out 3 times to obtain a meaningful number of reads spanning the entire LacO array.

For YBB32, YBB36, and YBB41, raw reads from the ONT runs were *de novo* assembled using Flye (v. 2.9.2) in nano-hq mode while specifying the genome length as 14 Mb. For YBB41, reads from the 3 sequencing attempts were first concatenated before *de novo* assembly. For YBB44, the Plasmidsaurus *de novo* assembly was used. Reads were mapped back to the appropriate *de novo* assemblies using Minimap2 (v. 2.28-r1209) (*66, 67*), ensuring assembly contiguity. Samtools (v. 1.6) (*68*) was used to filter out secondary and supplementary alignments, sort remaining primary aligned reads, and index aligned reads. The resulting alignment was visualized in the Integrated Genomics Viewer (v. 2.13.2) (*69*). We noted that the average read depth varied along YAC-*Hi*-4q21^LacO^. For instance, the 4q21 region has higher read depth than that of the *Hi* stuffer DNA. To ensure that portions of the BAC harboring 4q21 had not integrated elsewhere into YAC-*Hi* or the natural yeast genome, reads mapping to 4q21 were extracted and then Blast aligned to the natural yeast genome and YAC-*Hi*, revealing an absence of hits. We conclude that the most likely cause of this degree of variation is restricted to rearrangements and amplification within the sequences present in the BAC that harbors 4q21^LacO^. As well, we conclude that the salient design elements (gene expression cassette, LacO array, and 4q21 sequence to house centromeric chromatin) are present within YAC-*Hi*-4q21^LacO^. The YBB41 contig representing chromosome 8 was missing ∼26 kb, most of which was due to a collapse of the *CUP1* locus that organizes as a tandem array (*70*), and further exhibited poor coverage over deleted regions and their junctions, suggesting misassembly. A new *de novo* assembled genome was thus built with the YBB36 chromosome 8 contig. Alignment back to this contig resolved contiguity, consistent with misassembly of the YBB41 dataset. Note that *de novo* assembly for all strains failed to resolve chromosome 12, containing the rDNA locus, as a single contig, as previously reported (*8*). As a result, lengths depicted for chromosome 12 in genome representations throughout (Fig. 1D, 3B, and fig. S2A) are derived from the Saccer3 genome. The remaining chromosome lengths are direct outputs of *de novo* assembly.

To evaluate LacO array length, Seqtk (v. 1.4-r122) (https://github.com/lh3/seqtk) was used to retrieve pass-filter reads mapping to 300 unique nucleotides upstream and downstream of the LacO array. The length of each LacO array, considered the distance between the first bp of the first monomer and the last bp of the last monomer, was divided by the number of bp per monomer (39.85, including the monomer and variable-length intervening sequence) to generate the number of LacO copies for each read.

### Spheroplast fusion with HT1080^Dox-inducible mCherry-LacI-HJURP^ and U2OS^Dox-inducible mCherry-LacI-HJURP^ cells

YAC-*Hi*-4q21^LacO^ yeast were grown in synthetic minimal media (-histidine-uracil-tryptophan) to OD_600_ = 0.8-1.0 before harvesting at 1741 x g for 3 min. Yeast were resuspended in 20 ml 1 M sorbitol and incubated overnight at 4 ℃. Yeast were harvested at 1741 x g for 3 min and resuspended in 20 ml SPE medium. 40 µl β-mercaptoethanol and 100 µl 10 mg/ml zymolyase-20T were added to the yeast solution before gently inverting and then incubating at 37 ℃ in a shaker set to 25 rpm for 1 h. 30 ml ice cold 1 M sorbitol was then added to the spheroplasts before pelleting at 627 x g, 18 min, 4 ℃. The spheroplasts were washed in 50 ml ice cold 1 M sorbitol before harvesting at 627 x g, 25 min, 4 ℃. The pellet was resuspended in 1 ml STC, and the solution was allowed to incubate at room temperature for 10-30 min before addition to human cells.

Human cells were processed along with the spheroplasts. ∼4 h before yeast were pelleted, cells (∼60-80% confluent) were fed with fresh DMEM containing 10% certified tetracycline-free fetal bovine serum (FBS; Takara catalog #: 631106), 100 U/ml penicillin, and 100 µg/ml streptomycin (hereafter referred to as complete DMEM) and 50 µM S-trityl-L-cysteine (STLC) (Fisher: AAL1438406). After 5-6 h incubation in 50 µM STLC, cells were washed with PBS, trypsinized, and counted using a hemacytometer. Cells were pelleted and resuspended in PBS to a concentration of 6 x 10^5^ cells/ml. 3 x 10^5^ human cells were mixed with 9 x 10^7^ YAC-*Hi*-4q21^LacO^ spheroplasts and incubated at room temperature for 5 min. The mixture was pelleted at 1700 x g for 30 s in a microcentrifuge. The supernatant was aspirated and the pellet resuspended in a 45% PEG2000, 10% DMSO solution in 75 mM HEPES pH 7.5. The suspension was incubated at room temperature for 5 min. The fusion was quenched by adding 1 ml serum/antibiotic-free DMEM. Fused cells were pelleted at 1700 x g for 30 s in a microcentrifuge. Cells were resuspended in 1 ml complete DMEM and 500 µl aliquoted into two wells of a 6-well plate, each containing 2 ml complete DMEM and one containing a coverslip (for screening fusions). After 4 h (when cells have adhered), media is swapped with 3 ml complete DMEM +/- 2 µg/ml doxycycline (- doxycycline for coverslip wells intended for screening fusions). After 16 h, cells are again fed with 3 ml complete DMEM +/- 2 µg/ml doxycycline (- doxycycline for coverslip wells intended for screening fusions). Cells are allowed to incubate for 32 h before washing with PBS, trypsinizing, and transferred to a 10-cm plate with complete DMEM + G418 (325 µg/ml for HT1080^Dox-inducible mCherry-LacI-HJURP^ and 267 µg/ml for U2OS^Dox-inducible mCherry-LacI-HJURP^). Selection is maintained for 8 days before reducing selection to a maintenance concentration (150 µg/ml for HT1080^Dox-inducible^ ^mCherry-LacI-HJURP^ and 100 µg/ml for U2OS^Dox-inducible^ ^mCherry-LacI-HJURP^) for subsequent expansion. Note that wells containing coverslips are incubated for 72 h total before fixing, permeabilizing, and 4’-6-diamidino-2-phenylindole (DAPI) (100 ng/ml) staining. Fusion rates were ascertained from the fraction of cells harboring cytoplasmic yeast-derived, fluorescent punctae (fig. S6).

### IF-FISH on metaphase spreads

70-80% confluent human cells were cultured with complete DMEM containing 50 µM STLC and incubated at 37 ℃ for 2-4 h. Mitotic cells were blown off using a transfer pipet and pelleted at 415 x g for 5 min. Cells were resuspended in 500 µl PBS before counting with a hemacytometer. The cells were again pelleted at 415 x g for 5 min before dropwise resuspending the cells in hypotonic buffer consisting of a 1:1:1 ratio of 75 mM KCl, 0.8% Na_3_C_6_H_5_O_7_, and 1.5 mM MgCl_2_ to a concentration of 5 x 10^5^ cells/ml. Cells were allowed to swell at room temperature for 15 min before diluting to 5 x 10^4^ cells/ml in 500 µl total hypotonic buffer. Cells were cytospun at 1500 rpm, acceleration 9, for 5 min in a Shandon Cytospin 4 centrifuge. Cells were allowed to dry for 1 min before permeabilizing with KCM buffer (10 mM Tris-Cl pH 7.7, 120 mM KCl, 20 mM NaCl, and 0.1% Triton X-100) for 15 min. Spreads were blocked in IF block for 20 min before incubating with ⍺CENP-A (ENZO catalog #: ADI-KAM-CC006) diluted 1:1000 in IF block for 45 min at room temperature. Spreads were washed 3 times in KCM for 5 min each before incubating in Cy3-conjugated donkey anti-mouse IgG (Jackson Immunoresearch catalog #: 712-165-151) diluted to 3.5 µg/ml in IF block for 25 min. Spreads were again washed 3 times in KCM for 5 min before fixing in 4% paraformaldehyde for 10 min at room temperature. Spreads were washed 3 times with Milli-Q H_2_O before proceeding with FISH.

FISH probe was prepared by nick translation (Roche catalog #: 10976776001) using biotin-16-dUTP (Millipore catalog #: 11093070910) and a plasmid carrying the 10 kb LacO array (*26, 71*) as template. Unincorporated nucleotides were removed using Amersham MicroSpin G-50 columns (Cytiva catalog #: 27533002). 300 ng of probe/slide was ethanol precipitated along with 20 µg of salmon sperm DNA (Invitrogen catalog #: 15632011) and 10 µg of human Cot1 DNA (Invitrogen catalog #: 15279011). Precipitated DNA was resuspended in 50% formamide, 10% dextran sulfate in 2X SSC and denatured at 78 ℃ for 10 min before being placed in 37 ℃ for at least 20 min for pre-hybridization. 300 ng of probe was added to the RNase-treated spreads, and hybridization was performed at 37 ℃ for 16 h. Slides were washed 2 times in 50% formamide, 2X SSC at 37 ℃ for 5 min before washing slides 2 times in 2X SSC at 37 ℃ for 5 min. Slides were blocked with 2.5% milk in 4X SSC + 0.1% Tween 20 for 10 min and then incubated in NeutrAvidin-FITC (ThermoFisher catalog #: 31006) diluted to 25 µg/ml with 2.5% milk in 4X SSC + 0.1% Tween 20 at 37 ℃ for 1 h. Spreads were washed 3 times in 4X SSC + 0.1% Tween 20 at 45 ℃ at 45 ℃ for 2 min, followed by 1 wash in PBS. Spreads were then incubated in PBS supplemented with 1 µg/ml DAPI for 10 min at room temperature, washed again in PBS, and then water before mounting with Vectashield. Slides were imaged on an inverted epifluorescence microscope (Leica DMI6000B) with a 100x oil-immersion objective and the Leica K8 Scientific CMOS camera. Images were blind deconvolved within LAS X (v. 3.6.0.20104) while resizing from 12-bit depth to 16-bit depth. Channel contrasting and image cropping was carried out in ImageJ (v. 1.52k) before assembling figures in Adobe Illustrator (v. 29.7.1). The “HAC” designation was given to objects with FISH signal < 2 µm in diameter and 2 co-localized CENP-A foci.

### Visualizing HA-LacI-HJURP in recipient U2OS cells

YAC-*Hi*-4q21^LacO^ and YAC-*Hi*-4q21^LacO^; *pTPI1>HA-LacI-HJURP* spheroplasts were prepared as described above. The fusion was performed with U2OS cells (U2OS derivatives (*72*) that are the direct precursor cell line to U2OS^Dox-inducible^ ^mCherry-LacI-HJURP^ cells, described above). The fusion differed in the following fashion: complete DMEM was comprised of FBS (Sigma catalog #: F2442), 100 U/ml penicillin, and 100 µg/ml streptomycin; the cells were not cultured in STLC for 6 h prior to fusion but rather were cultured normally until harvesting by trypsinization. Post fusion, cells were cultured in the same complete DMEM without G418. 500 µl of the fused cell suspension was aliquoted into 2 wells of a 6-well plate containing glass coverslips. After 4 h, media was swapped on all wells. 20 h post fusion, the first timepoint of samples was collected and fixed in 4% paraformaldehyde. These coverslips were subsequently washed 2 times with 2 ml PBS and then stored at 4 ℃ in PBS. Media was swapped on the 72 h samples. At 72 h post fusion, the 72 h samples were fixed in 4% paraformaldehyde. All samples were quenched with 100 mM Tris-Cl (pH 7.5) for 5 min. Cells were permeabilized with PBS + 0.5% Triton X-100 for 5 min. Samples were washed 3 times with PBS + 0.1% Tween 20 (PBST) for 5 min each. Coverslips were blocked with 100 µl IF block (2% FBS, 2% BSA, 0.1% Tween 20, and 0.02% NaN_3_) for 20 min by floating coverslips before incubating with anti-HA antibody (Cell Signaling Technologies catalog #:C29F4 lot #: 10) diluted 1:1000 in IF block for 45 min at room temperature. Samples were washed 3 times with PBST for 5 min each. Coverslips were floated on FITC-conjugated goat anti-rabbit (Jackson Immunoresearch catalog #: 111-095-144) for 25 min at room temperature. Samples were washed 1 time in PBST for 5 min. Samples were DAPI stained with 100 ng/ml DAPI diluted in PBST for 5 min. Samples were washed 1 time in PBST for 5 min, then 1 time in PBS for 5 min, then left in Milli-Q H_2_O until mounting. Samples were mounted with 10 µl Vectashield. Imaging was performed with the same apparatus as described in “IF-FISH on metaphase spreads” but with a 40x oil-immersion objective.

Representative images were prepared post deconvolution as described in “IF-FISH on metaphase spreads.” Quantification was carried out on raw images in ImageJ (v. 1.52k). To do so, stacks of fields were max projected. Selections of individual cells (or groups of cells if segmentation was not possible) were measured and integrated density collected. Background was subtracted by averaging pixel intensities of three regions not covered by cells and multiplying by the selection area and subtracting this value from the integrated density of the selected cell(s). Integrated densities of background-subtracted selections were summed and then divided by the number of cells for which data was collected to derive the mean integrated density for the experiment. To account for signal that cannot be attributed to LacI-HJURP (i.e., from autofluorescence, non-specific antibody staining, or spectral overlap from tdTomato expression; for an example, see Fig. S5), the timepoint-matched average value for the YAC-*Hi*-4q21^LacO^ fusion experiments was subtracted from the mean integrated density of each YAC-*Hi*-4q21^LacO^; *pTPI1>HA-LacI-HJURP* fusion experiment. Note that timepoint-matched correction is necessary because signal that cannot be associated to LacI-HJURP accumulates in cells from 20 h to 72 h. Data were then normalized to the difference in the average fluorescence value of the YAC-*Hi*-4q21^LacO^; *pTPI1>HA-LacI-HJURP* fusion experiments and the YAC-*Hi*-4q21^LacO^ fusion experiments at 20 h for graphical display.

### HAC formation assay without antibiotic selection in U2OS cells

YAC-*Hi*-4q21^LacO^ and YAC-*Hi*-4q21^LacO^; *pTPI1>HA-LacI-HJURP* spheroplasts were prepared as described above. The assay differed in the following fashion. Complete DMEM was comprised of FBS (Sigma catalog #: F2442), 100 U/ml penicillin, and 100 µg/ml streptomycin. Interphase cells were harvested by trypsinization. After spheroplast fusion, cells were seeded in complete DMEM, which was refreshed after 4 h and again after 16 h. Cells were expanded to 10-cm plates and either cryogenically stored or subjected to HAC formation experimentation.

### IF-FISH on interphase cells

Spheroplasts and U2OS cells were prepared as described in “Visualizing HA-LacI-HJURP in recipient U2OS cells” above. 250 µl U2OS cell suspensions after spheroplast fusion were seeded directly into the middle of a circle drawn by a hydrophobic pen on slides coated with poly-D-lysine (Thermo Fisher catalog #: 67-762-15). 250 µl of fresh complete DMEM was then added to the circle, and the slides were incubated at 37 ℃ in a closed 15 cm plate containing an open 6 cm plate filled with sterile H_2_O for humidity. After 12 h, slides were washed with PBS, fixed for 10 min in 4% paraformaldehyde, washed 2 times in PBS, and stored in a humidified box at 4 ℃ in 500 µl PBS. After 72 h, relevant slides were washed with PBS and fixed with 4% paraformaldehyde. All samples were quenched together with 100 mM Tris-Cl (pH 7.5). Cells were permeabilized with KCM for 15 min and then washed 3 times in KCM for 3 min. IF was performed as described in “IF-FISH on metaphase spreads” but with affinity-purified anti-CENP-C rabbit polyclonal antibody diluted to 0.85 µg/ml in IF block (for detecting centromere formation on YACs) or with anti-⍺-tubulin (Sigma-Aldrich catalog #: T9026) diluted to 10.6 µg/ml in IF block (for defining the U2OS cell area). Cy3-conjugated goat anti-rabbit IgG (Jackson Immunoresearch catalog #: 111-165-144) diluted to 3.5 µg/ml was used as secondary in CENP-C detection. Cy3-conjugated donkey anti-mouse IgG (Jackson Immunoresearch catalog #: 712-165-151) diluted to 3.5 µg/ml was used as secondary for ⍺-tubulin detection. FISH was performed as described in “IF-FISH on metaphase spreads.” For measuring yeast nuclei in cells after fusion, FISH was performed using nick-translated probes prepared from YBB24 (see Table S1) HMW gDNA extracted as described in “Yeast strain sequencing, genome assembly, and analysis” and further purified using the Qiaquick PCR purification kit. For identifying YACs, FISH was performed using nick-translated probes prepared from the plasmid carrying the 10 kb LacO array. Images were acquired and processed as described in “IF-FISH on metaphase spreads.” FISH signal spanning 0.9-2 µm were used to score yeast genomes. YACs were considered CENP-C+ if signal was above background and LacO and CENP-C foci were within 200 nm of one another.

### Sorting single cells harboring HACS via flow cytometry

U2OS cells derived from the non-selection HAC formation assay (Fig. 5, A to C) were grown to ∼80% confluence before washing with PBS and trypsinizing. Cells were pelleted at 300 x g for 5 min at room temperature and washed 2 times with 10 ml PBS containing 1 mM EDTA. Cells were again harvested at 300 x g for 5 min at room temperature before resuspending in 1 ml PBS containing 1 mM EDTA and 1% BSA. The cells were placed in 5 ml sterile polysterene tubes and stored on ice until sorting. Sorting was performed at the Flow Cytometry Core Laboratory at The Children’s Hospital of Philadelphia on the FACSAria Fusion using a 100 µm nozzle. U2OS cells were first gated on SSC-A x FSC-A, then gated for singlets using FSC-H x FSC-A. For the -enrichment by fluorescence sort, one singlet was deposited per well. For the +enrichment by fluorescence sort, an additional SSC-A x PE-A gate was applied, and one tdTomato+ singlet (fig. S7) was deposited per well. Wells contained 150 µl 50% conditioned media derived from parental U2OS cells. Briefly, U2OS cells were grown to 40% confluence before replacing their media. Cells were then grown overnight, media harvested, spun down at 300 x g for 5 min at room temperature, and filtered through a 0.22 µm filter. 100% conditioned media was then mixed 1:1 with fresh complete media and used within a week.

Plates of cells were incubated at 37 ℃ for 2-3 weeks before wells containing successfully grown colonies were passaged to a well of a 24-well plate. Once ∼80% confluent, cells were passaged to a well of a 6-well plate. Once 6-well plates were ∼80% confluent, 83% of the cell suspension was cryogenically stored, and the remaining 17% was passaged to a 6-cm plate. Once these cells were ∼80% confluent, chromosome spreads were prepared and CENP-A IF, LacO FISH performed (see “IF-FISH on metaphase spreads” for details). In total, cells were grown for ∼4 weeks before IF-FISH was performed.

As monoclonal HAC lines were derived in the absence of selective pressure, we quantified stability as the minimum retention rate, discriminating it from the conventional approach for measuring HAC retention, which is normally performed on clones derived in the presence of drug selection (*1, 3, 6*).

Since the starting HAC fraction of a HAC clone must be a single cell, retention rate can be inferred because an ∼80% confluent 6-cm plate of U2OS cells has ∼1.5 x 10^6^ cells. As cells were diluted 1:6 into the 6-cm plate, an appropriate estimate for the total number of cells grown is 9 x 10^6^, which occurred thr<u>ough a minimum of 2</u>3 cell doublings. We can therefore calculate the minimum retention rate as 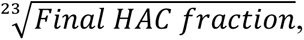 noting “minimum” as the cell growth estimation does not take cell death into account, which would increase the true number of cell doublings between clonal isolation and IF-FISH. Further, early mis-segregation events would exacerbate perception of stability relative to its true stability.

---

**Figure S1:**
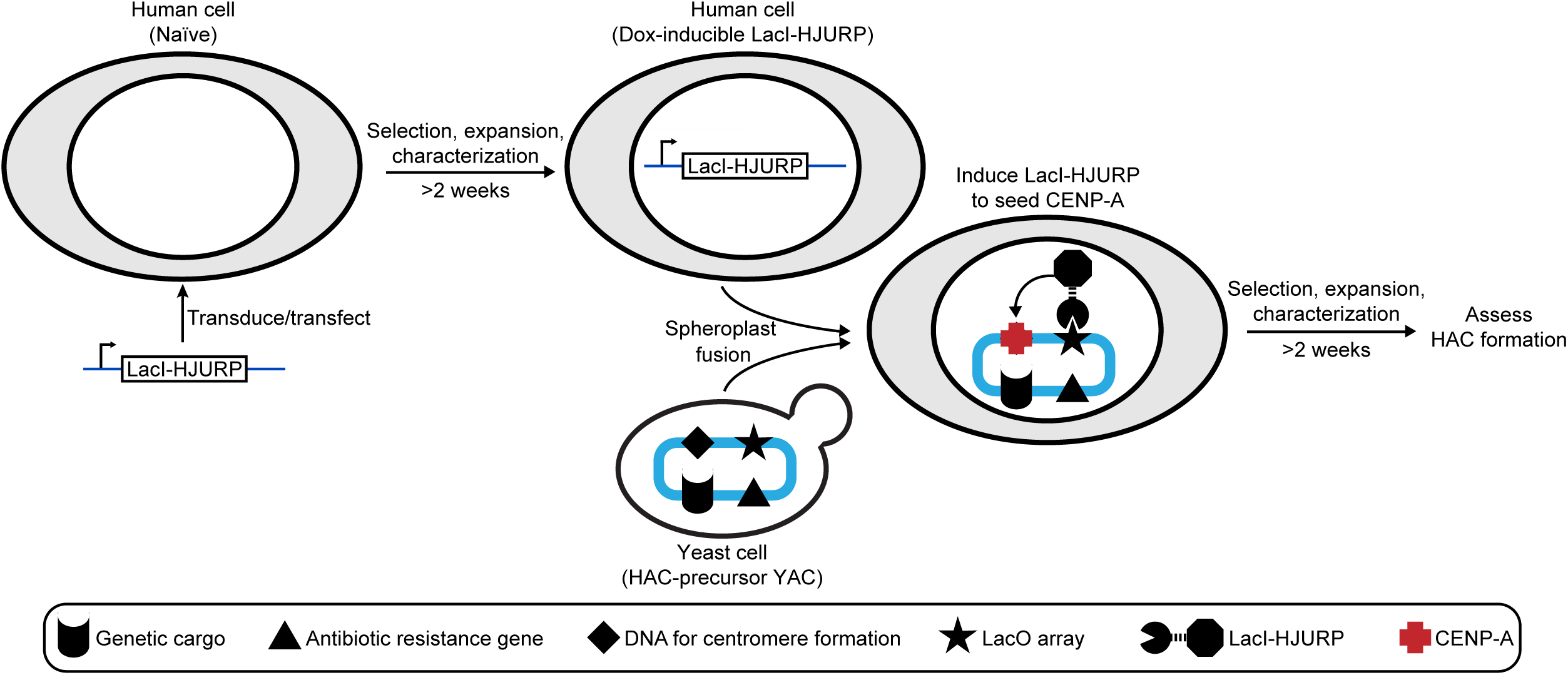
Process to genetically engineer HAC formation competence and form HACs. Candidate human cells for HAC formation are first transfected/transduced to introduce a dox-inducible LacI-HJURP expression cassette. Antibiotic selection is performed to derive the HAC formation-competent cell line, which is subsequently expanded and characterized. These cells are then fused with yeast to receive the HAC-precursor YAC, and centromere formation is induced by a 48-h doxycycline treatment. Antibiotic selection is again performed to enrich for stable events including HACs, and that cell population is then subjected to quantification of HAC formation events.

**Figure S2:**
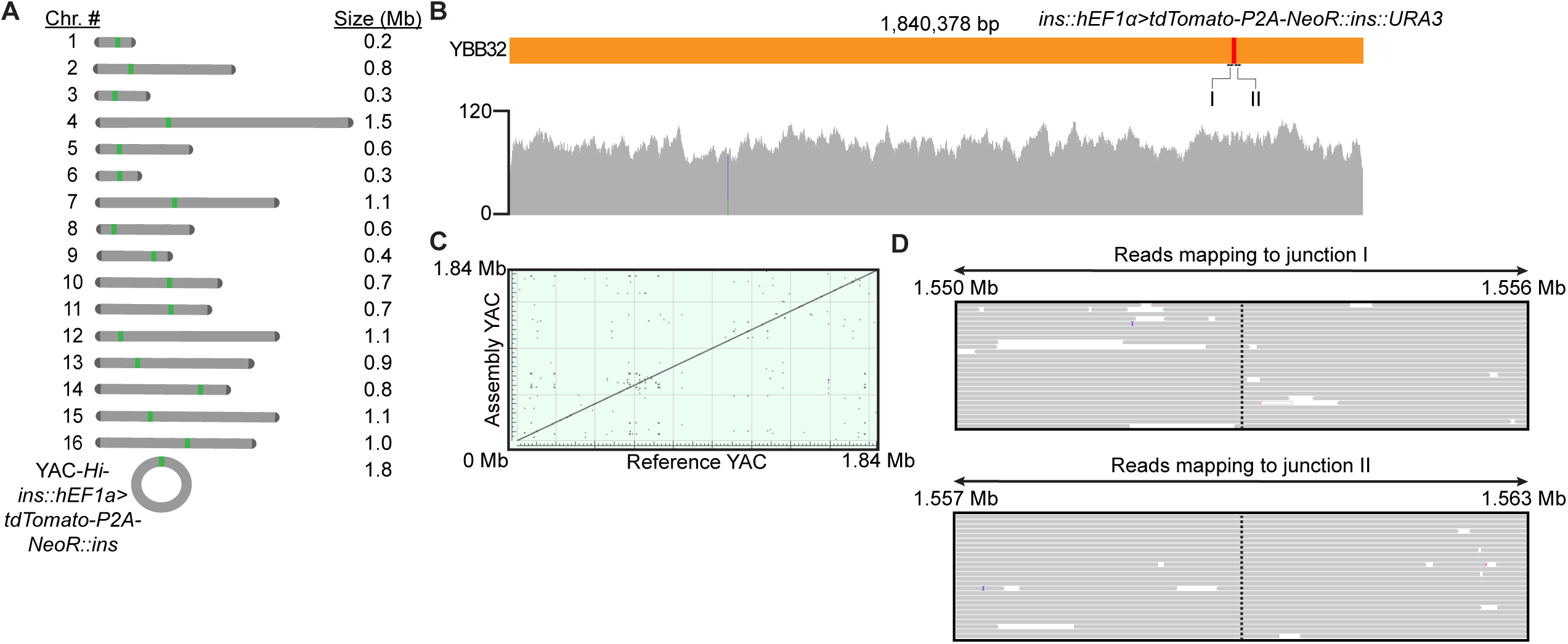
Generating and characterizing YAC-*Hi*-ins::*hEF1⍺>tdTomato-P2A-NeoR*::ins yeast. (A) Representation of the yeast chromosomes derived from ONT sequencing and an unbiased assembly of pass filter reads. (B) Integrated Genomics Viewer alignment track of reads mapped to the *de novo* assembled YAC. (C) Dot plot generated by BLAST alignment of the *de novo* assembled YAC-*Hi*-ins::*hEF1⍺>tdTomato-P2A-NeoR*::ins, referred to as “Assembly YAC,” against the predicted recombinant YAC, referred to as “Reference YAC.” (D) Reads spanning a 6 kb region that includes the *H. flu*-expression cassette junction and a 6 kb region that includes the expression cassette-*H. flu* junction. The vertical purple bar indicates a nucleotide insertion in the read relative to the mapped reference sequence.

**Figure S3:**
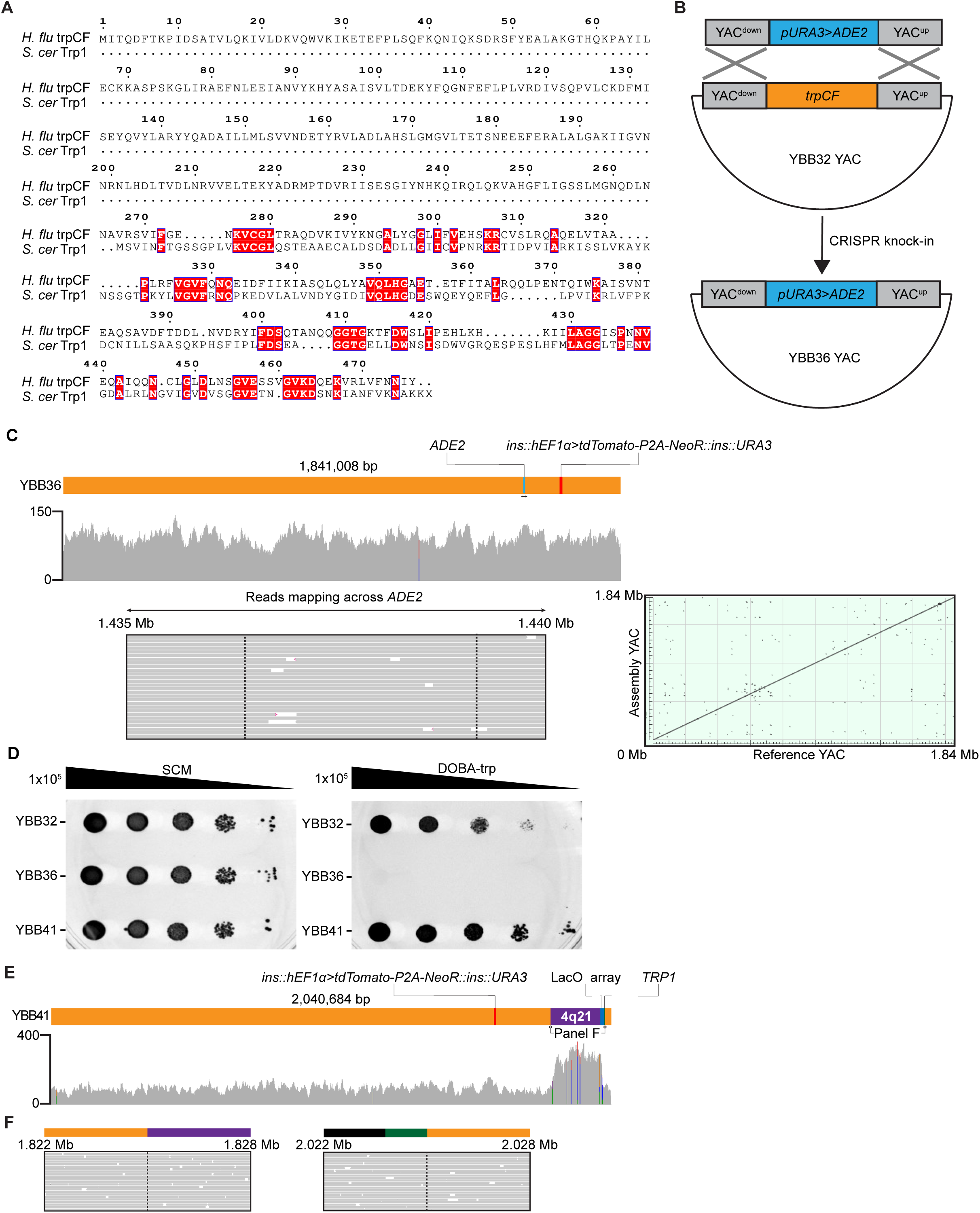
Deleting *trpCF* restores utility of *TRP1* as an auxotrophic marker to engineer YAC-*Hi*-4q21^LacO^ yeast strain via TAR, and contiguous alignment track and reads spanning 4q21^LacO^ integration site reveal on-target modification and no alterations of the remaining backbone. (A) *S. cerevisiae* Trp1 was BLAST aligned to *Hi* trpCF to produce a .aln file. This file was then input into ESPript 3.0 to generate the multiple sequence alignment. The (phosphoribosyl)anthranilate isomerase (trpF) domain spans residues 276-476 of the annotated multifunctional polypeptide and shares 28% identity with *S. cerevisiae* Trp1. (B) Recombination strategy to replace *trpCF* with an *ADE2* cassette, enabling red-white screening of recombinant strains. The cassette is flanked by 300-bp homology arms. (C) Integrated Genomics Viewer alignment track of reads mapped to the *de novo* assembled YAC. Reads mapping across the *ADE2* integration are shown with junctions demarcated by dotted lines. A dot plot of the *de novo* assembled YAC, termed “Assembly YAC,” aligned against the *in silico* prediction, termed “Reference YAC.” (D) Spot assay screening -tryptophan prototrophy of each of YBB32(*trpCF+*; *TRP1-)*, YBB36(*trpCF-*; *TRP1-*), and YBB41(*trpCF-*; *TRP1+*). 1 x 10^5^ yeast cells were spotted in the left column, and then 10-fold serial dilutions were spotted down each of the right columns. Yeast spotted on synthetic complete medium (SCM) serves as a control for the number of plated yeast. (E) Integrated Genomics Viewer alignment track of reads to the *de novo* assembled YAC-*Hi*-4q21^LacO^. Grey denotes a perfect match, blue denotes the unexpected presence of a cytosine, red denotes the unexpected presence of a thymine, copper denotes the unexpected presence of a guanine, and green denotes the unexpected presence of an adenine. Note that the increase in coverage throughout 4q21 can be attributed to concatemerization of regions within the BAC during integration. Importantly, reads mapping to 4q21 do not map to the natural yeast genome or off-target locations of the *Hi* stuffer (see **Materials and Methods** for additional details), suggesting that concatemerization is confined to the on-target integration and that all design elements (mammalian expression cassette, LacO array, and 4q21) are present within YAC-*Hi*-4q21^LacO^. (F) Reads seamlessly spanning a 6 kb region that includes the *H. flu*-4q21 junction and a 6 kb region that includes the *TRP1*-*H. flu* junction reveals on-target integration.

**Figure S4:**
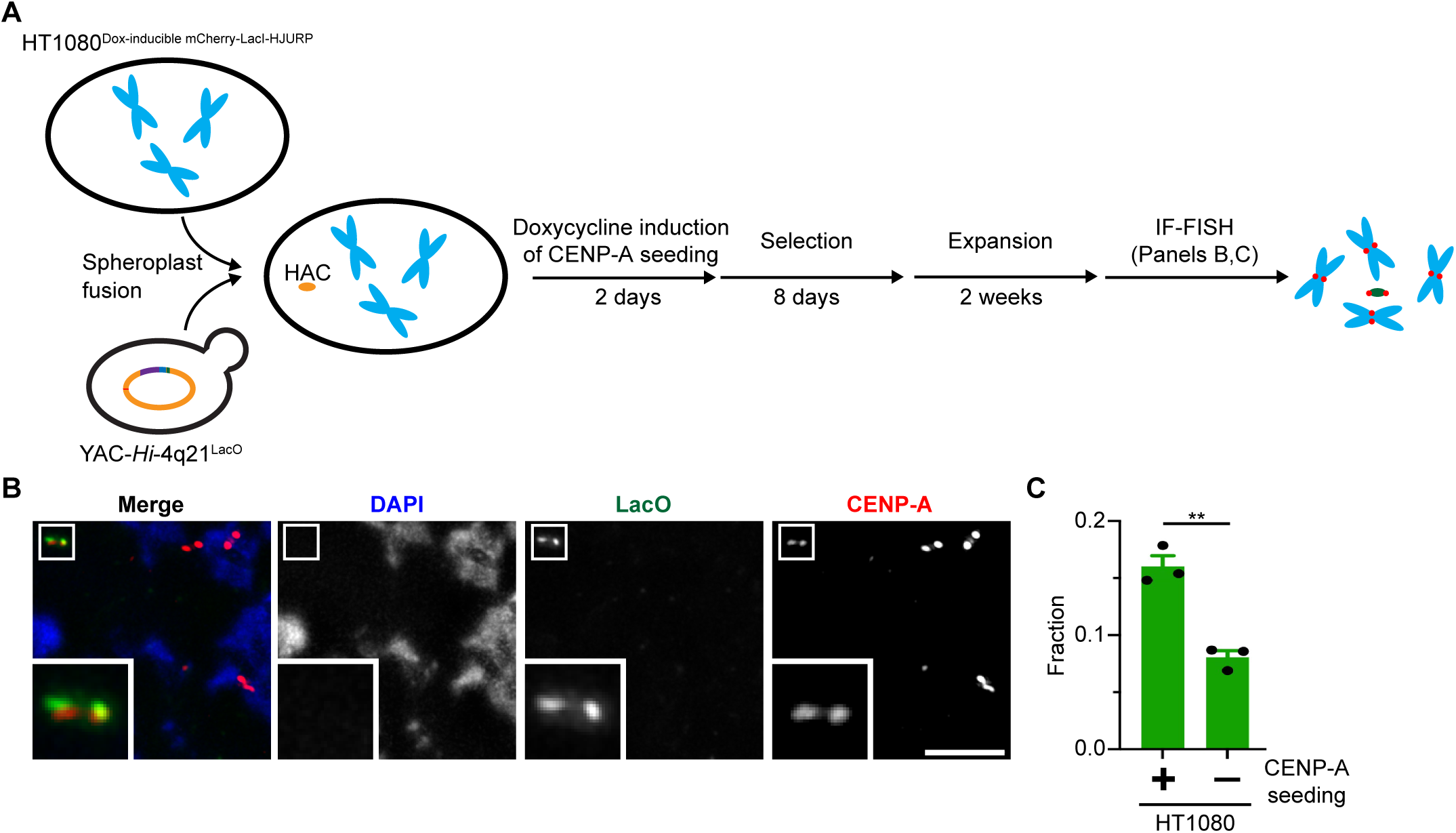
HAC formation by fusion of HT1080^Dox-inducible^ ^mCherry-LacI-HJURP^ cells with spheroplasts carrying YAC-*Hi*-4q21^LacO^. (A) Graphical depiction of experiment. (B) Representative image of HACs formed in HT1080^Dox-inducible^ ^mCherry-LacI-HJURP^ cells treated with doxycycline following fusion with YAC-*Hi*-4q21^LacO^ spheroplasts. Bar, 5 µm. Inset, 5X magnification. (C) Quantification of HAC formation frequency +/- doxycycline-mediated induction of mCherry-LacI-HJURP expression, revealing a similar rate (16.0 +/- 0.9%) of HACs formed after dox induction as exhibited by the U2OS^Dox-inducible^ ^mCherry-LacI-HJURP^ cells (Figure 2). The mean (+/- standard error) is shown. The increase in HAC frequency was assessed using an unpaired, one-tailed t test. ** denotes p < 0.01, n=3 (≥ 20 cells/replicate).

**Figure S5:**
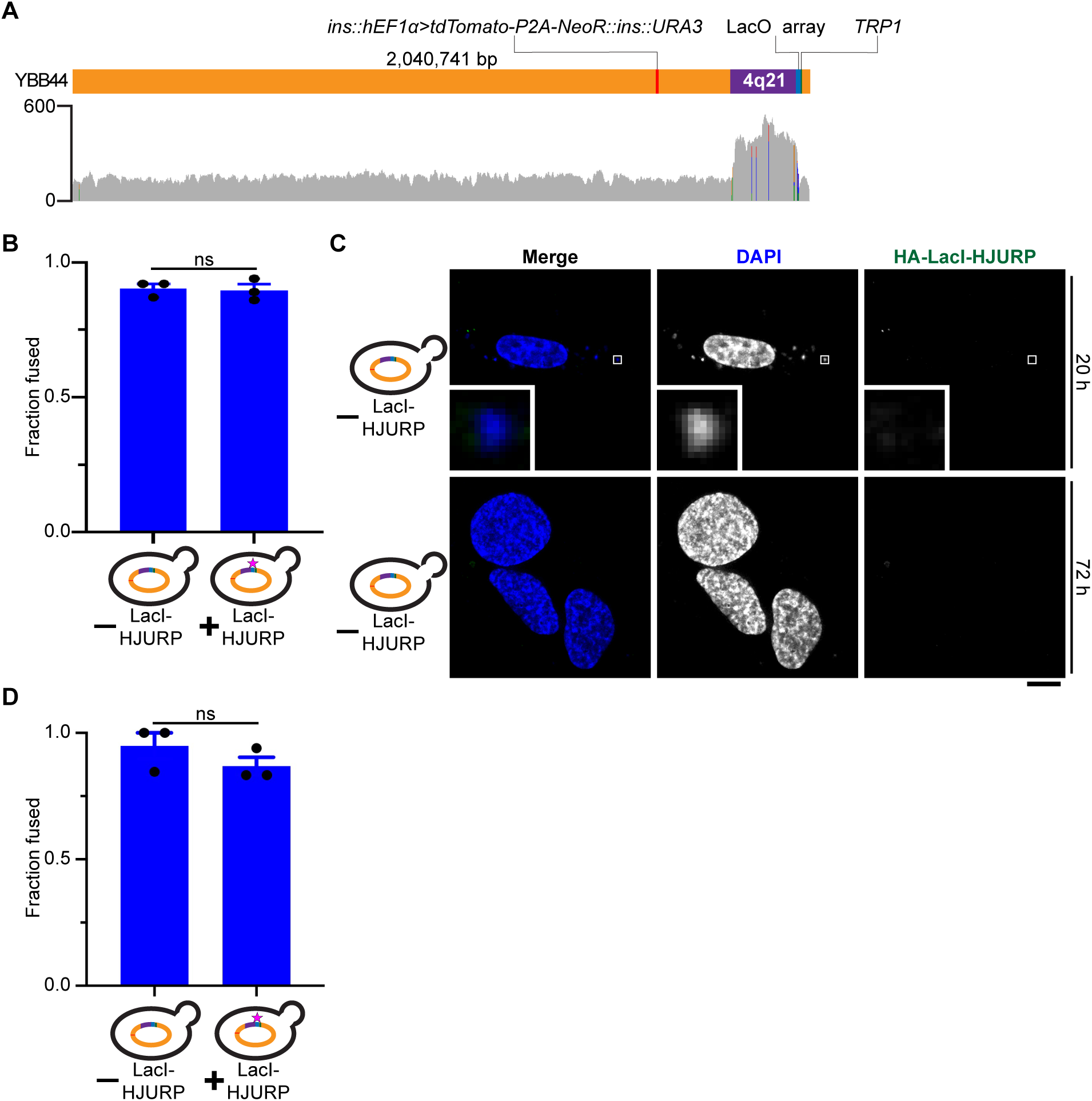
YAC-*Hi*-4q21^LacO^ remained stable through introduction of *pTPI1>HA-LacI-HJURP* into natural yeast chromosome 4 and controls for assessing early molecular events following fusion of U2OS cells with YAC-*Hi*-4q21^LacO^; *pTPI1>HA-LacI-HJURP* spheroplasts. (A) Integrated Genomics Viewer alignment track of reads mapped back to the *de novo* assembled YAC. Coverage and depth of coverage are similar to that of the YAC-*Hi*-4q21^LacO^ strain, suggesting stable maintenance of the YAC through transformation and outgrowth. (B) Quantification of spheroplast fusion efficiency in experiment to assess LacI-HJURP delivery (see Fig. 3, F to H) as dictated by the presence of yeast nuclei observed in the cytoplasm of U2OS cells 20 h post fusion. The difference in fusion rate between YAC-*Hi*-4q21^LacO^ and YAC-*Hi*-4q21^LacO^; *pTPI1>HA-LacI-HJURP* spheroplast fusions was evaluated with an unpaired, two-tailed t test. P > 0.05, n=3 (≥ 22 cells/replicate). (C) Representative images of U2OS cells fused with YAC-*Hi*-4q21^LacO^ spheroplasts and stained for HA-LacI-HJURP 20 h and 72 h post fusion. Fused U2OS cells at 20 h exhibit cytoplasmic yeast nuclei and little non-specific IF signal. By 72 h, most of the cytoplasmic yeast nuclei are absent and IF signal is again non-specific and/or autofluorescence associated with the fusion. Bar, 10 µm. Inset, 10X magnification. (D) Quantification of spheroplast fusion efficiency in experiment to assess early centromere formation (see Fig. 4, A to C) as dictated by the presence of yeast nuclei observed in the cytoplasm of U2OS cells 12 h post fusion. The difference in fusion rate between YAC-*Hi*-4q21^LacO^ and YAC-*Hi*-4q21^LacO^; *pTPI1>HA-LacI-HJURP* spheroplast fusions was evaluated with an unpaired, two-tailed t test. P > 0.05, n=3 (≥ 26 cells/replicate).

**Figure S6:**
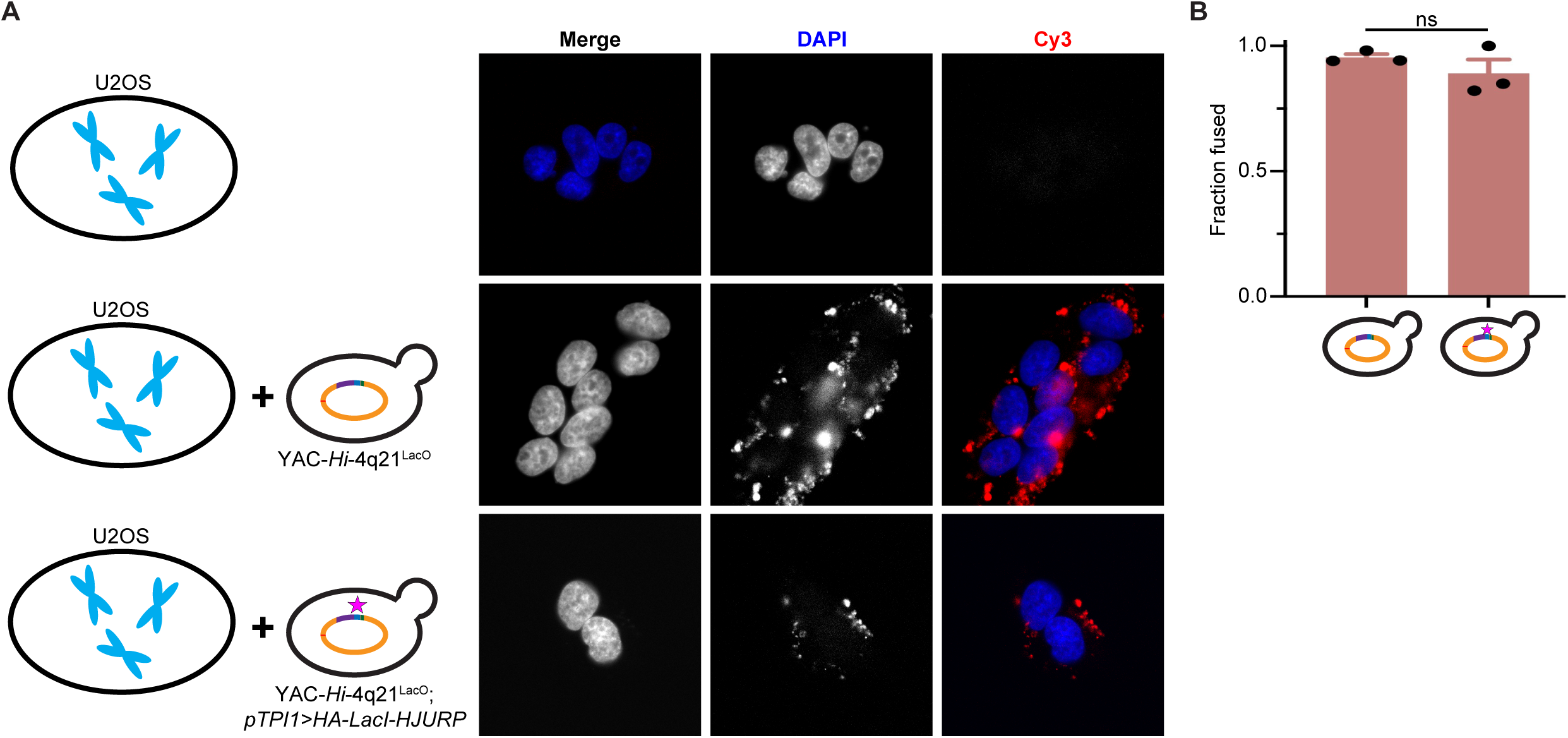
Efficiency of spheroplast fusion with U2OS cells for non-selection HAC formation assay. (A) Representative fluorescence images of cells that were treated with fusion buffer, mixed with YAC-*Hi*-4q21^LacO^ spheroplasts before treating with fusion buffer, or mixed with YAC-*Hi*-4q21^LacO^; *pTPI1>HA-LacI-HJURP* spheroplasts before treating with fusion buffer. Note that much of the cytoplasmic focal signal can be attributed to autofluorescence of spheroplast material present in the cytoplasm readily visualized with a filter appropriate for Cy3 fluorescence. (B) Quantification of spheroplast fusion efficiency as dictated by the presence of yeast-derived cytoplasmic foci. The difference in fusion rate between YAC-*Hi*-4q21^LacO^ and YAC-*Hi*-4q21^LacO^; *pTPI1>HA-LacI-HJURP* spheroplasts was evaluated with an unpaired, two-tailed t test. P > 0.05, n=3 (≥ 50 cells/replicate).

**Figure S7:**
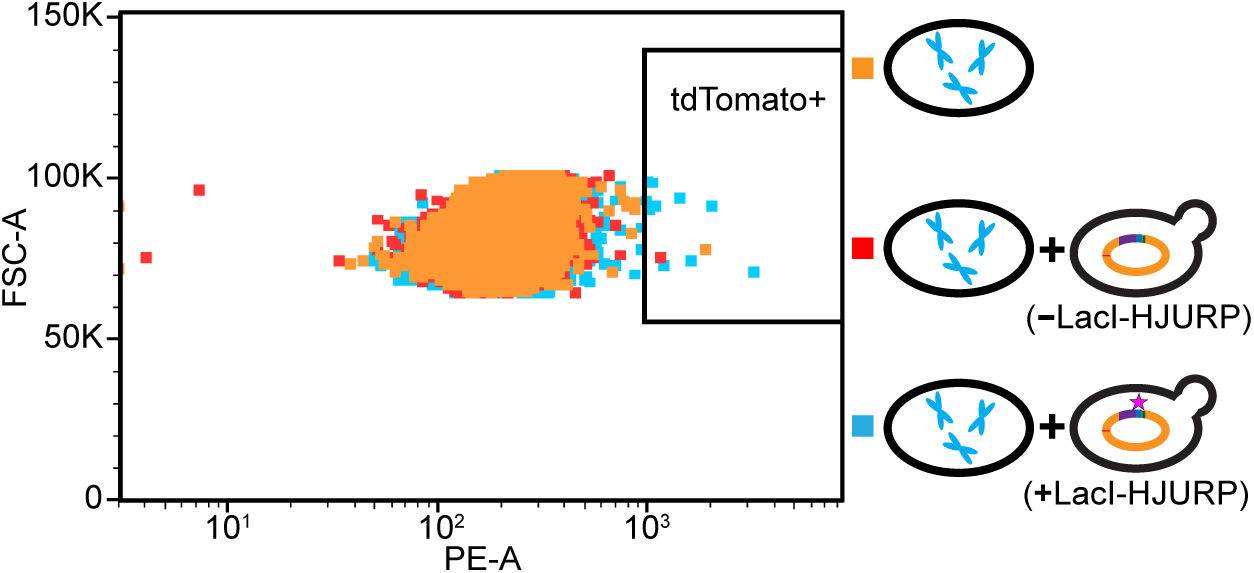
Strategy to derive monoclonal HAC lines with or without gating for cell fluorescence attributable to tdTomato. U2OS cells were subjected to the fusion protocol (with/without the indicated yeast strains) and subcultured for ∼4 weeks before sorting. Shown is the proportion (0.09%) of U2OS cells fused to YAC-*Hi*-4q21^LacO^; *pTPI1>HA-LacI-HJURP* spheroplasts that are tdTomato+. We interpret the tdTomato+ fraction to indicate gene silencing with rare escape permissive to the fluorescence and associated HAC clone enrichment we observe (see Fig. 5D).

**Table S1:** Yeast strains. Yeast strains used in this study.

| Strain | Description | Source |
| --- | --- | --- |
| YBB24 | VL6-48N yeast | Reference 73 |
| YBB28 | VL6-48 yeast with YAC-*Hi* | Reference 41 |
| YBB31 | YBB28 with Cas9 YCp | This study |
| YBB32 | YBB31 with ins::*hEF1α*>*tdTomato-P2A-NeoR*::ins integrated into YAC-*Hi* by TAR | This study |
| YBB36 | YBB32 with *trpCF* in YAC-*Hi* replaced with *ADE2* | This study |
| YBB41 | YBB36 with 4q21<sup>LacO</sup> integrated into YAC-*Hi* by TAR | This study |
| YBB44 | YBB41 with *pTPI1*>*HA-LacI-HJURP* integrated into natural chromosome 4 | This study |

**Table S2:**
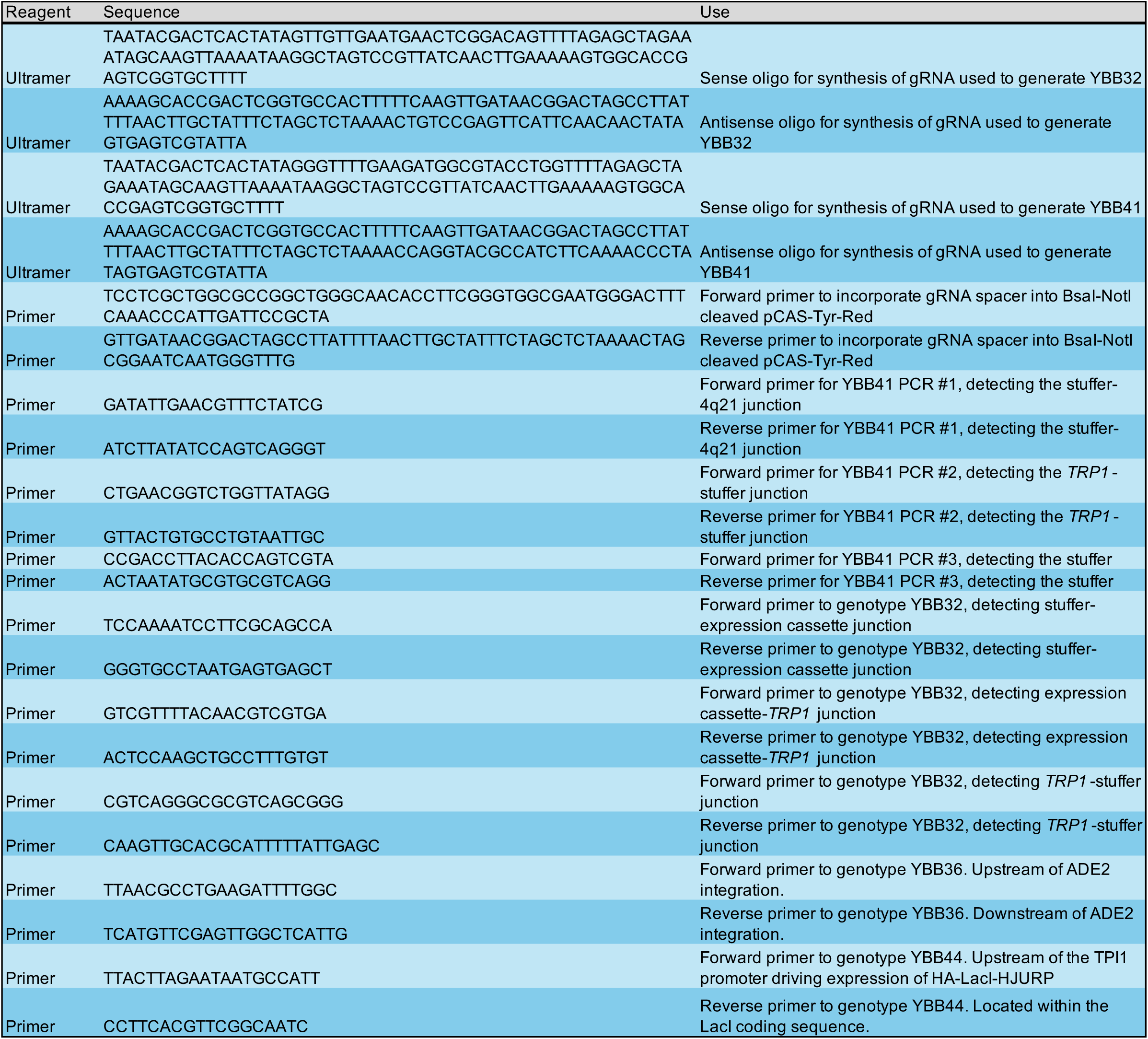
Oligonucleotides. Primer and Ultramer sequences used in this study.

## References

1. J. J. Harrington, G. V. Bokkelen, R. W. Mays, K. Gustashaw, H. F. Willard, Formation of de novo centromeres and construction of first-generation human artificial microchromosomes. Nature Genetics 15, 345–355 (1997).

2. M. Ikeno et al., Construction of YAC–based mammalian artificial chromosomes. Nature Biotechnology 16, 431–439 (1998).

3. T. A. Ebersole et al., Mammalian artificial chromosome formation from circular alphoid input DNA does not require telomere repeats. Hum Mol Genet 9, 1623–1631 (2000).

4. M. K. Rudd, R. W. Mays, S. Schwartz, H. F. Willard, Human artificial chromosomes with alpha satellite-based de novo centromeres show increased frequency of nondisjunction and anaphase lag. Mol Cell Biol 23, 7689–7697 (2003).

5. M. G. Schueler, A. W. Higgins, M. K. Rudd, K. Gustashaw, H. F. Willard, Genomic and genetic definition of a functional human centromere. Science 294, 109–115 (2001).

6. G. A. Logsdon et al., Human artificial chromosomes that bypass centromeric DNA. Cell 178, 624–639 (2019).

7. C. W. Gambogi et al., Efficient formation of single-copy human artificial chromosomes. Science 383, 1344–1349 (2024).

8. G. J. Birchak et al., Rapid assembly of functional modules for generating human artificial chromosome constructs compatible with epigenetic centromere seeding. Chromosome Res 33, 29 (2025).

9. N. Annaluru et al., Total synthesis of a functional designer eukaryotic chromosome. Science 344, 55–58 (2014).

10. B. A. Blount et al., Synthetic yeast chromosome XI design provides a testbed for the study of extrachromosomal circular DNA dynamics. Cell Genom 3, 100418 (2023).

11. J. S. Dymond et al., Synthetic chromosome arms function in yeast and generate phenotypic diversity by design. Nature 477, 471–476 (2011).

12. C. A. Hutchison et al., Design and synthesis of a minimal bacterial genome. Science 351, 1414–1425 (2016).

13. W. E. Robertson et al., Escherichia coli with a 57-codon genetic code. Science 390, 351–364 (2025).

14. D. Schindler et al., Design, construction, and functional characterization of a tRNA neochromosome in yeast. Cell 186, 5237–5253 (2023).

15. Y. Zhao et al., Debugging and consolidating multiple synthetic chromosomes reveals combinatorial genetic interactions. Cell 186, 5220–5236 (2023).

16. J. S. James, J. Dai, W. L. Chew, Y. Cai, The design and engineering of synthetic genomes. Nat Rev Genet 26, 298–319 (2025).

17. D. B. Thompson et al., The future of multiplexed eukaryotic genome engineering. ACS Chem Biol 13, 313–325 (2018).

18. J. D. Boeke et al., The Genome Project–Write. Science 353, 126–127 (2016).

19. D. Schindler, R. S. K. Walker, Y. Cai, Methodological advances enabled by the construction of a synthetic yeast genome. Cell Reports Methods 4, 100761 (2024).

20. L. Clarke, J. Carbon, Isolation of a yeast centromere and construction of functional small circular chromosomes. Nature 287, 504–509 (1980).

21. K. Kixmoeller, P. K. Allu, B. E. Black, The centromere comes into focus: from CENP-A nucleosomes to kinetochore connections with the spindle. Open Biol 10, 200051 (2020).

22. A. Musacchio, A. Desai, A molecular view of kinetochore assembly and function. Biology 6, 5 (2017).

23. W. C. Earnshaw, N. Rothfield, Identification of a family of human centromere proteins using autoimmune sera from patients with scleroderma. Chromosoma 91, 313–321 (1985).

24. D. K. Palmer, K. O’Day, M. H. Wener, B. S. Andrews, R. L. Margolis, A 17-kD centromere protein (CENP-A) copurifies with nucleosome core particles and with histones. J Cell Biol 104, 805–815 (1987).

25. M. C. Barnhart et al., HJURP is a CENP-A chromatin assembly factor sufficient to form a functional de novo kinetochore. J Cell Biol 194, 229–243 (2011).

26. M. J. Mendiburo, J. Padeken, S. Fülöp, A. Schepers, P. Heun, Drosophila CENH3 is sufficient for centromere formation. Science 334, 686–690 (2011).

27. T. Hori, W. H. Shang, K. Takeuchi, T. Fukagawa, The CCAN recruits CENP-A to the centromere and forms the structural core for kinetochore assembly. J Cell Biol 200, 45–60 (2013).

28. C. C. Chen et al., CAL1 is the Drosophila CENP-A assembly factor. J Cell Biol 204, 313–329 (2014).

29. G. A. Logsdon et al., Both tails and the centromere targeting domain of CENP-A are required for centromere establishment. J Cell Biol 208, 521–531 (2015).

30. A. Guse, C. W. Carroll, B. Moree, C. J. Fuller, A. F. Straight, In vitro centromere and kinetochore assembly on defined chromatin templates. Nature 477, 354–358 (2011).

31. B. Moree, C. B. Meyer, C. J. Fuller, A. F. Straight, CENP-C recruits M18BP1 to centromeres to promote CENP-A chromatin assembly. J Cell Biol 194, 855–871 (2011).

32. D. R. Foltz et al., Centromere-specific assembly of CENP-A nucleosomes is mediated by HJURP. Cell 137, 472–484 (2009).

33. E. M. Dunleavy et al., HJURP is a cell-cycle-dependent maintenance and deposition factor of CENP-A at centromeres. Cell 137, 485–497 (2009).

34. M. C. Silva et al., Cdk activity couples epigenetic centromere inheritance to cell cycle progression. Dev Cell 22, 52–63 (2012).

35. S. Muller et al., Phosphorylation and DNA binding of HJURP determine its centromeric recruitment and function in CenH3 (CENP-A) loading. Cell Rep 8, 190–203 (2014).

36. D. Conti et al., Role of protein kinase PLK1 in the epigenetic maintenance of centromeres. Science 385, 1091–1097 (2024).

37. D. G. Gibson et al., Creation of a bacterial cell controlled by a chemically synthesized genome. Science 329, 52–56 (2010).

38. W. Zhang et al., Manipulating the 3D organization of the largest synthetic yeast chromosome. Mol Cell 83, 4424–4437 (2023).

39. R. K. Dawe, Engineering better artificial chromosomes. Science 383, 1292–1293 (2024).

40. D. G. Gibson et al., Complete chemical synthesis, assembly, and cloning of a Mycoplasma genitalium genome. Science 319, 1215–1220 (2008).

41. B. J. Karas et al., Direct transfer of whole genomes from bacteria to yeast. Nat Methods 10, 410–412 (2013).

42. C. Lartigue et al., Creating bacterial strains from genomes that have been cloned and engineered in yeast. Science 325, 1693–1696 (2009).

43. L. Meneu et al., Sequence-dependent activity and compartmentalization of foreign DNA in a eukaryotic nucleus. Science 387, eadm9466 (2025).

44. D. M. C. Bittencourt et al., Minimal bacterial cell JCVI-syn3B as a chassis to investigate interactions between bacteria and mammalian cells. ACS Synth Biol 13, 1128–1141 (2024).

45. J. K. Keesey, Jr, J. A. Demoss, Cloning of the trp-J gene from Neurospora crassa by complementation of a trpC mutation in Escherichia coli. J Bacteriol 152, 954–958 (1982).

46. N. Kouprina, V. Larionov, Selective isolation of genomic loci from complex genomes by transformation-associated recombination cloning in the yeast Saccharomyces cerevisiae. Nat Protoc 3, 371–377 (2008).

47. D. L. Neil et al., Structural instability of human tandemly repeated DNA sequences cloned in yeast artificial chromosome vectors. Nucleic Acids Res 18, 1421–1428 (1990).

48. T. Fukagawa, W. R. A. Brown, Efficient conditional mutation of the vertebrate CENP-C gene. Human Molecular Genetics 6, 2301–2308 (1997).

49. H. Kato et al., A conserved mechanism for centromeric nucleosome recognition by centromere protein CENP-C. Science 340, 1110–1113 (2013).

50. P. K. Allu et al., Structure of the human core centromeric nucleosome complex. Curr Biol 29, 2625–2639 (2019).

51. D. M. Brown et al., Efficient size-independent chromosome delivery from yeast to cultured cell lines. Nucleic Acids Res 45, e50 (2017).

52. J. T. D. Dunnen et al., Reconstruction of the 2.4 Mb human DMD-gene by homologous YAC recombination. Hum Mol Genet 1, 19–28 (1992).

53. Y. Shao et al., Creating a functional single-chromosome yeast. Nature 560, 331–335 (2018).

54. S. Nurk et al., The complete sequence of a human genome. Science 376, 44–53 (2022).

55. J. K. Jadlowsky et al., Long-term safety of lentiviral or gammaretroviral gene-modified T cell therapies. Nat Med 31, 1134–1144 (2025).

56. J. Maher, R. J. Brentjens, G. Gunset, I. Rivière, M. Sadelain, Human T-lymphocyte cytotoxicity and proliferation directed by a single chimeric TCRζ/CD28 receptor. Nat Biotechnol 20, 70–75 (2002).

57. G. Ferrari, A. J. Thrasher, A. Aiuti, Gene therapy using haematopoietic stem and progenitor cells. Nat Rev Genet 22, 216–234 (2021).

58. M. S. Singh et al., Retinal stem cell transplantation: Balancing safety and potential. Prog Retin Eye Res 75, 100779 (2020).

59. C. Sun, C. Serra, G. Lee, K. R. Wagner, Stem cell-based therapies for Duchenne muscular dystrophy. Exp Neurol 323, 113086 (2020).

60. J. S. Lanni, T. Jacks, Characterization of the p53-dependent postmitotic checkpoint following spindle disruption. Mol Cell Biol 18, 1055–1064 (1998).

61. J. D. Orth, A. Loewer, G. Lahav, T. J. Mitchison, Prolonged mitotic arrest triggers partial activation of apoptosis, resulting in DNA damage and p53 induction. Mol Biol Cell 23, 567–576 (2012).

62. D. G. Gibson, Gene and genome construction in yeast. Curr Protoc Mol Biol Chapter 3, Unit3 22 (2011).

63. B. D. M. Bean, M. Whiteway, V. J. J. Martin, The MyLO CRISPR-Cas9 toolkit: a markerless yeast localization and overexpression CRISPR-Cas9 toolkit. G3 (Bethesda) 12, (2022).

64. R. D. Gietz, Yeast transformation by the LiAc/SS carrier DNA/PEG method. Methods Mol Biol 1205, 1–12 (2014).

65. K. Kannan et al., One step engineering of the small-subunit ribosomal RNA using CRISPR/Cas9. Sci Rep 6, 30714 (2016).

66. H. Li, Minimap2: pairwise alignment for nucleotide sequences. Bioinformatics 34, 3094–3100 (2018).

67. H. Li, New strategies to improve minimap2 alignment accuracy. Bioinformatics 37, 4572–4574 (2021).

68. H. Li et al., The Sequence Alignment/Map format and SAMtools. Bioinformatics 25, 2078–2079 (2009).

69. J. T. Robinson et al., Integrative genomics viewer. Nat Biotechnol 29, 24–26 (2011).

70. S. Fogel, J. W. Welch, G. Cathala, M. Karin, Gene amplification in yeast: CUP1 copy number regulates copper resistance. Curr Genet 7, 347–355 (1983).

71. C. C. Robinett et al., In vivo localization of DNA sequences and visualization of large-scale chromatin organization using Lac operator/repressor recognition. J Cell Biol 135, 1685–1700 (1996).

72. P. Khandelia, K. Yap, E. V. Makeyev, Streamlined platform for short hairpin RNA interference and transgenesis in cultured mammalian cells. Proc Natl Acad Sci U S A 108, 12799–12804 (2011).

73. V. Noskov et al., A genetic system for direct selection of gene-positive clones during recombinational cloning in yeast. Nucleic Acids Res 30, E8 (2002).

